# Multi-scale spatial genetic structure of a vector-borne plant pathogen in orchards and wild habitat

**DOI:** 10.1101/795096

**Authors:** Véronique Marie-Jeanne, François Bonnot, Gaël Thébaud, Jean Peccoud, Gérard Labonne, Nicolas Sauvion

## Abstract

Inferring the dispersal processes of vector-borne plant pathogens is a great challenge because the plausible epidemiological scenarios often involve complex spread patterns at multiple scales. European stone fruit yellows (ESFY), a disease caused by ‘*Candidatus* Phytoplasma prunorum’ and disseminated via planting material and vectors belonging to the species *Cacopsylla pruni*, is a major threat for stone fruit production throughout Europe. The spatial genetic structure of the pathogen was investigated at multiple scales by the application of a combination of statistical approaches to a large dataset obtained through the intensive sampling of the three ecological compartments hosting the pathogen (psyllids, wild and cultivated *Prunus*) in three *Prunus*-growing regions in France. This work revealed new haplotypes of ‘*Ca*. P. prunorum’, and showed that the prevalence of the different haplotypes of this pathogen is highly uneven between all regions, and within two of them. In addition, we identified a significant clustering of similar haplotypes within a radius of at most 50 km, but not between nearby wild and cultivated *Prunus*. We also provide evidence that the two species of the *C. pruni* complex are unevenly distributed but can spread the pathogen, and that infected plants are transferred between production areas. Altogether, this work supports a main epidemiological scenario where ‘*Ca*. P. prunorum’ is endemic in, and mostly acquired from, wild *Prunus* by immature *C. pruni* (of both species) who then migrate to “shelter plants” that epidemiologically connect sites less than 50 km apart by later providing infectious mature *C. pruni* to their “migration basins”, which differ in their haplotypic composition. We argue that such multiscale studies would be very useful for other pathosystems.

## INTRODUCTION

Vector-borne pathogens have caused some of the most devastating plant diseases in perennial and annual crops^1,2^. Managing these threats requires understanding their epidemiological cycle, in particular the dispersal processes of the pathogens in the landscape^3^. However, deciphering the complexity of the plausible epidemiological scenarios poses a great challenge, especially as insects are involved in disease spread. Indeed, almost all vector-borne plant pathogens are transmitted by small piercing-sucking insects of the Hemiptera order, which are known to disperse at various distances while searching their host plants for food resources and/or reproduction^4^. Added to this question of dispersal distance is the issue of different host plant species that insects can occupy during their life cycle and their movements among them. In this context, a key question that often arises is the role of wild plants as a reservoir for the pathogen and/or the insect vector^5–10^.

One of the first difficulties in identifying patterns of disease spread in agricultural landscapes is to determine the optimal scale of the study design. Although the spatial scale of the processes has been a central question for decades in ecology^11–14^ and population genetics^15,16^, the development of large-scale pattern-oriented approaches to understand the processes shaping the genetic structure of a population is recent^17–21^. The basic idea of these approaches is to estimate the distance at which two samples become genetically independent by relating genetic data and spatial information obtained for a set of samples, through combining methods of geostatistics^22^ and population or landscape genetics^23^. In particular, approaches used for qualitative data comprise join-counts^24^ and permutation tests to identify distances at which genetic dissimilarity between pairs of individuals is significantly lower than expected^25–28^.

The present study focuses on a disease of fruit trees named European stone fruit yellows (ESFY), caused by ‘*Candidatus* Phytoplasma prunorum’^29^, and disseminated via planting material^30^ or the psyllid *Cacopsylla pruni* (Scopoli, 1763)^31^. The phytoplasma and its psyllid vectors are widespread in Europe, including the most important stone fruit production areas, where the disease significantly impacts susceptible crops, in particular apricot and Japanese plum trees^30,32^. Rigorous sanitary control of nursery plants and insecticide treatments are currently the main disease-control strategies, but despite these measures ESFY continues to economically impact European fruit growers, which raises the question of the origin of contaminations.

Over the past 20 years, several studies have aimed to decipher the complexity of the phytoplasma/*Prunus*/psyllid pathosystem^33^. Two points appear essential in the dynamics of the epidemic: (i) in Europe, the pathogen is often found in wild *Prunu*s^32,34^, and (ii) the psyllid vectors are univoltine and migrate twice a year between *Prunus* and conifers^9^. In a landscape comprising different plants where the psyllid vectors can feed, different mutually nonexclusive scenarios of spread of the pathogen in the orchards can be proposed (Fig. 1). Thébaud et al.^9^ showed that the most likely way of orchard contamination involved mature (i.e. immigrant) psyllids carrying the phytoplasma at a sufficient concentration for inoculation (e.g., scenarios 3 and 4 in Fig. 1). Nevertheless, this lab-based conclusion has not yet been backed by field studies investigating the role played by local secondary spread of ESFY by immature (i.e. emigrant) or mature *C. pruni* involved in either within-orchard tree-to-tree transmission (e.g., scenario 1 in Fig. 1) or acquisition of the phytoplasma in bushes before transmission to a nearby orchard (e.g., scenario 2 in Fig. 1). The relative contribution of natural spread and human transfer of contaminated plants is another major unknown in this pathosystem (scenarios 7_a+b_ and 8 in Fig. 1). Finally, the spatial scale of these processes is unknown and, in combination with the landscape structure, this may lead to more or less complex epidemiological networks connecting close or distant ecological compartments (Supplementary Fig. S1).

**Figure 1.**
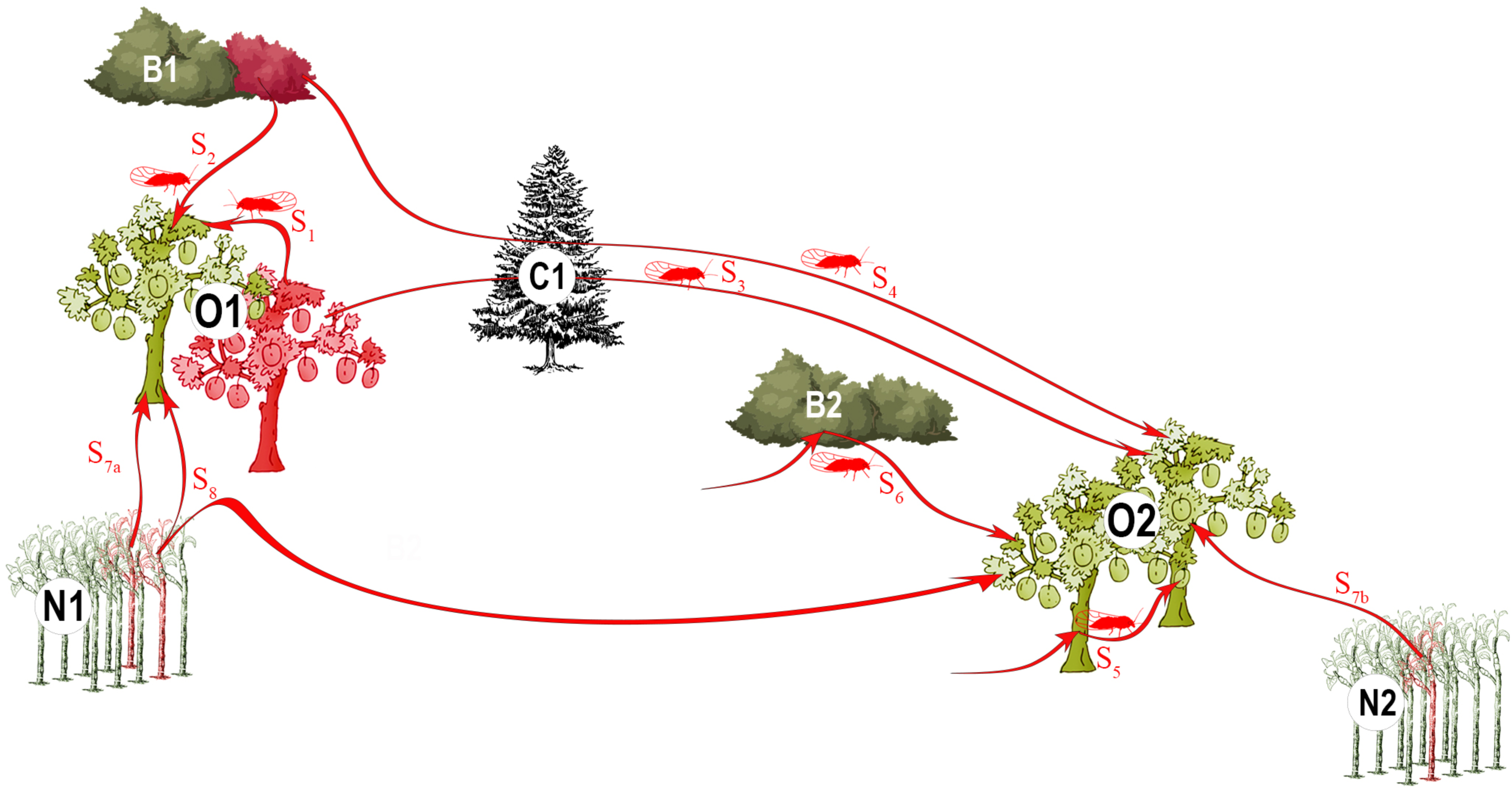
Eight mutually nonexclusive eco-epidemiological scenarios through which ‘*Ca*. P. prunorum’ might be spread in orchards. Bi: bushes (i.e. wild *Prunus* = host plants); Ci: conifers (i.e. shelter plants); Ni: nurseries; Oi: orchards. In red: cultivated trees, nursery plants, and wild *Prunus* infected by the phytoplasma, or infectious psyllids; in green: non-infected plants. Pathogen transmission to cultivated trees may involve various spatio-temporal scales depending on the involved scenario: transmission to a healthy apricot tree by a psyllid that acquired the pathogen from a nearby infected cultivated tree (S1) or a nearby infected bush (S2); transmission by a mature psyllid that acquired the phytoplasma on an infected tree (S3) or bush (S4) the previous year; multiple transmissions by the same infectious psyllid to nearby cultivated trees (S5), or to a bush and then a nearby cultivated tree (S6); independent contaminations of orchards by plants from nearby nurseries (S7a + S7b); contaminations of distant orchards by plants from the same nursery (S8).

The goal of the present work is to improve our understanding of the spatial scale and ecological compartments involved in ESFY epidemics, by investigating the spatial genetic structure of ‘*Ca*. P. prunorum’ using complementary statistical approaches combining genetic data and geographical distances. Our study was conducted in three French growing regions [Pyrénées-Orientales (PO), Bas-Rhône (BR) and Valence (VA)] that are strongly impacted by the disease, and was based on intensive sampling of the three main hosts of the pathogen (psyllids, wild and cultivated *Prunus*).

## RESULTS

### Genetic diversity of ‘*Ca*. Phytoplasma prunorum’

In the three sampling regions, we collected: (i) plant samples in infected orchards (mainly apricot), (ii) plant samples in wild *Prunus* bushes (mainly blackthorn) at various distances from these orchards, and (iii) mature psyllids in these bushes (Supplementary Figs. S2, S3, S4 and S5). A total of 6,342 samples were collected and molecularly tested for the presence of ‘*Ca*. P. prunorum’ (Table 1). The prevalence of ‘*Ca*. P. prunorum’ in bushes was high: on average 43.3% of the samples were found positive for the phytoplasma, from which we could obtain 339 sequences of the immunodominant membrane protein (*imp*) gene. From the 69 sampled orchards, 750 samples were found positive for the phytoplasma, and we obtained 553 sequences. In the fifteen orchards where a systematic sampling was undertaken, 14.3% of the 1,982 sampled trees were found positive for the phytoplasma (Supplementary Table S1), but only 4.3% of these were asymptomatic. Among 117 samples from symptomatic trees sampled several times (at different times or on different branches at the same date), six showed distinct *imp* haplotypes. Among the 2,572 psyllids collected from 71 different bushes, 104 (4.8%) were found carrying the phytoplasma, from which we obtained 99 sequences. Thus, 991 samples were successfully genotyped for the *imp* gene (Table 1).

**Table 1.** Statistics on the samples from each of the three ecological compartments (bush, psyllid, and orchard), all regions combined and within each region. PO: Pyrénées-Orientales; BR: Bas-Rhône; VA: Valence. n_1_: number of orchards or bushes per region, in which plant or insect samples were collected. n_2_: mean number of plant samples (or psyllids) collected per orchard or bush [minimum and maximum values]. N: total number of samples collected per region. #ESFY+: total number of samples found positive by PCR. %ESFY+: mean percentage of PCR-positive plant samples (or psyllids) per orchard or bush [minimum and maximum values]. #IMP: number of samples where the *imp* gene was successfully genotyped.

| Region | Compartment | n<sub>1</sub> | n<sub>2</sub><br>[min-max] | N | #ESFY+ | %ESFY+<br>[min-max] | #IMP |
| --- | --- | --- | --- | --- | --- | --- | --- |
| PO+BR+VA | bush | 61 | 21 [5-46] | 1114 | 474 | 43.3 [0-100] | 339 |
|  | psyllid | 71 | 34 [1-196] | 2572 | 104 | 4.8 [0-50.0] | 99 |
|  | orchard | 69 | 37 [1-244] | 2656 | 750 | 13.0 [1.9-39.3] | 553 |
| | $\Sigma$ | 201 | | 6342 | 1328 | | 991 |
| PO | bush | 14 | 24 [10-46] | 332 | 163 | 47.4 [0-100] | 123 |
|  | psyllid | 32 | 54 [7-196] | 1723 | 61 | 3.5 [0.7-14.3] | 58 |
|  | orchard | 34 | 28 [1-255] | 953 | 331 | 17.1 [1.9-39.3] | 267 |
| | $\Sigma$ | 80 | | 3008 | 555 | | 448 |
| BR | bush | 33 | 13 [5-40] | 426 | 152 | 39.2 [0-100] | 99 |
|  | psyllid | 11 | 27 [2-58] | 299 | 19 | 6.4 [1.9-50.0] | 19 |
|  | orchard | 24 | 38 [1-140] | 901 | 254 | 12.5 [4.0-25.9] | 164 |
| | $\Sigma$ | 68 | | 1626 | 425 | | 282 |
| VA | bush | 14 | 25 [16-33] | 356 | 159 | 43.4 [0-100] | 117 |
|  | psyllid | 28 | 20 [1-76] | 550 | 24 | 4.4 [0-50.0] | 22 |
|  | orchard | 11 | 44 [2-244] | 802 | 165 | 9.3 [2.7-19.3] | 122 |
| | $\Sigma$ | 53 | | 1708 | 348 | | 261 |

We identified 17 *imp* haplotypes, haplotypes I01 and I09 being the most frequent ones (44.2% and 30.0% of the samples, respectively; Supplementary Table S2). Four other *imp* haplotypes (i.e., I04, I10, I11, I13) reach also a frequency above 4%. Thus, only these six major haplotypes were considered in the subsequent statistical analyses. Among the eleven other haplotypes, I03 was already described in Germany by Danet et al.^35^ and I10-310 was recently described in Slovenia by Dermastia et al.^36^ under the name I34. Nine haplotypes (i.e., I01-104, I01-248, I01-339, I04-9, I04-175, I04-407, I04-453, I10-307, and I11-267) had not been described before (Supplementary Table S2). Conversely, five haplotypes (I02, I06, I07, I08, I12) previously described in France by Danet et al.^35^ were not found in our survey. The haplotype network illustrating the genetic relationships among the haplotypes shows that most of the obtained haplotypes differ by only one or two mutations, whereas haplotypes I13 and I05 differ widely from all other haplotypes (Fig. 2).

**Figure 2.**
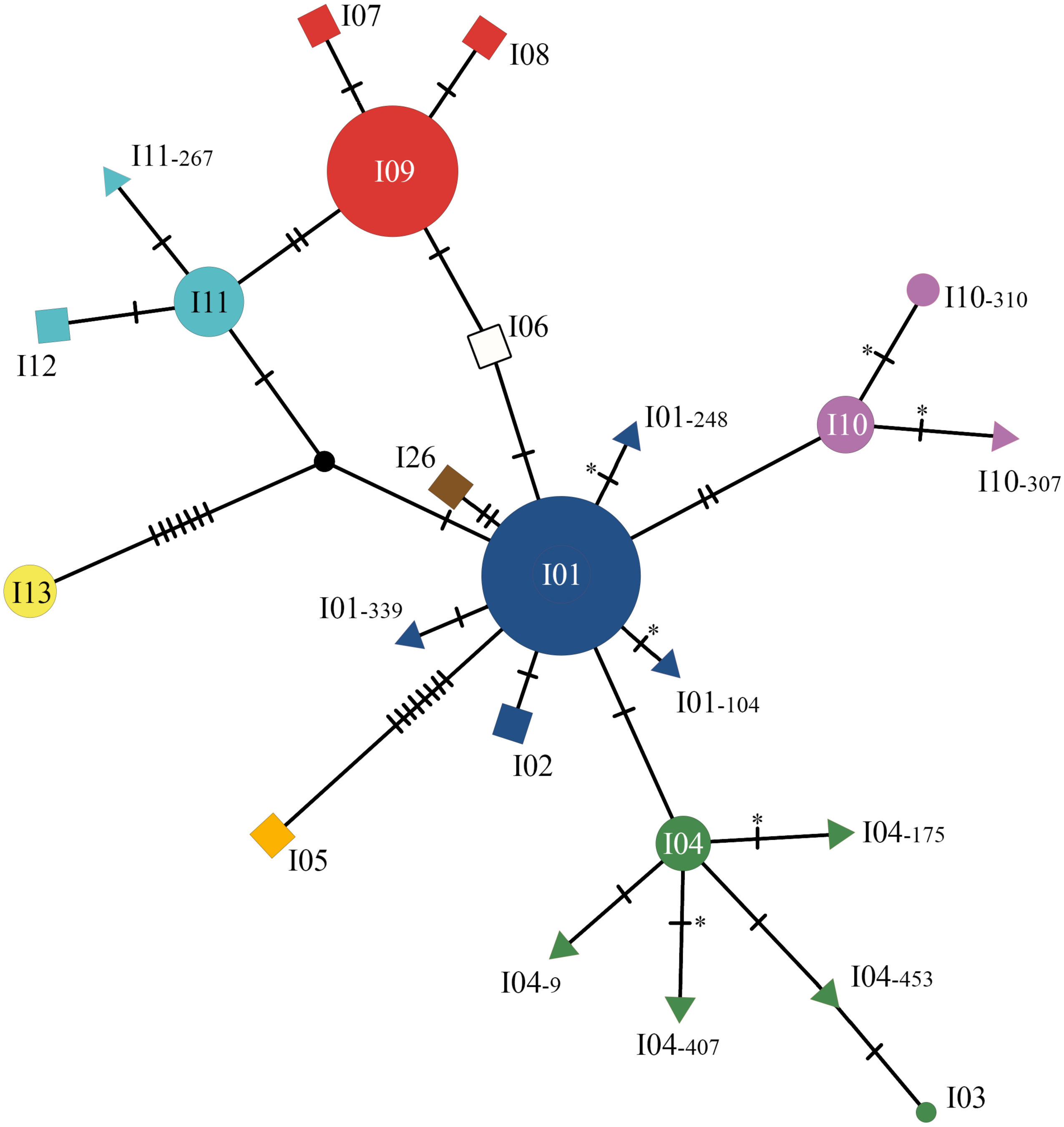
Haplotype network inferred from 24 *imp* sequences of ‘*Ca*. P. prunorum’ using the integer neighbour-joining algorithm. The disks represent the previously described haplotypes that were found in our samples. Disk area is proportional to the number of genotyped samples (Supplementary Table S2). The squares represent the previously described haplotypes that were not found in our samples. The triangles represent the newly described haplotypes. Each of them was named according to the major haplotype from which it derived, followed by the position of the mutation in the open reading frame of the *imp* DNA sequence. Hatch marks along edges represent the number of mutations differentiating two connected sequences. Asterisks indicate changes in the amino acid sequence. The black node represents an inferred unsampled sequence.

The psyllid vector *C. pruni* is known to present two cryptic species currently called A and B, which show clear genetic differences despite being ecologically and morphologically indistinguishable^37,38^. All the collected psyllids were successfully assigned to either species. Among all regions combined, species A was more frequent (62.6% of the samples; Supplementary Table S5), but the proportion varies greatly depending on the region, in particular in BR where this species strongly prevails (97.3%). There was no significant difference in phytoplasma prevalence (grey cells in Supplementary Table S5) between psyllid species across all regions (*p*=0.52; Fisher’s exact test). The proportions of the six major haplotypes differed significantly between the two species (*p*=2×10^−4^ across all regions; Fisher’s exact test), mainly because of the over-representation of haplotype I01 in species B. For the following analysis, the *C. pruni* complex was considered as a single epidemiological entity because performing distinct analyses for species A and B strongly reduced the statistical power.

### Exploratory data analysis

In order to explore relationships among haplotypes, compartments and regions, we performed a correspondence analysis on the contingency table of haplotype frequencies (Supplementary Table S3). The first two axes of the correspondence analysis explain a large part (78.7%) of the variance (Fig. 3), thus the obtained dimension reduction can be considered as very acceptable to interpret the data. The main modalities contributing to axis 1 are haplotype I09 and BR bushes and to a lesser extent haplotype I01, while the main modalities contributing to axis 2 are haplotype I11, VA orchards and PO bushes (Fig. 3b). In the PO region, all compartments appear very closely related, and well separated from those of the VA region. The BR region has an intermediate profile, and the three compartments were more dissimilar than for the other regions (Fig. 3a). In addition, the haplotypes from orchards were genetically more related (i.e., segregating only on axis 2) than those of the other two compartments.

**Figure 3.**
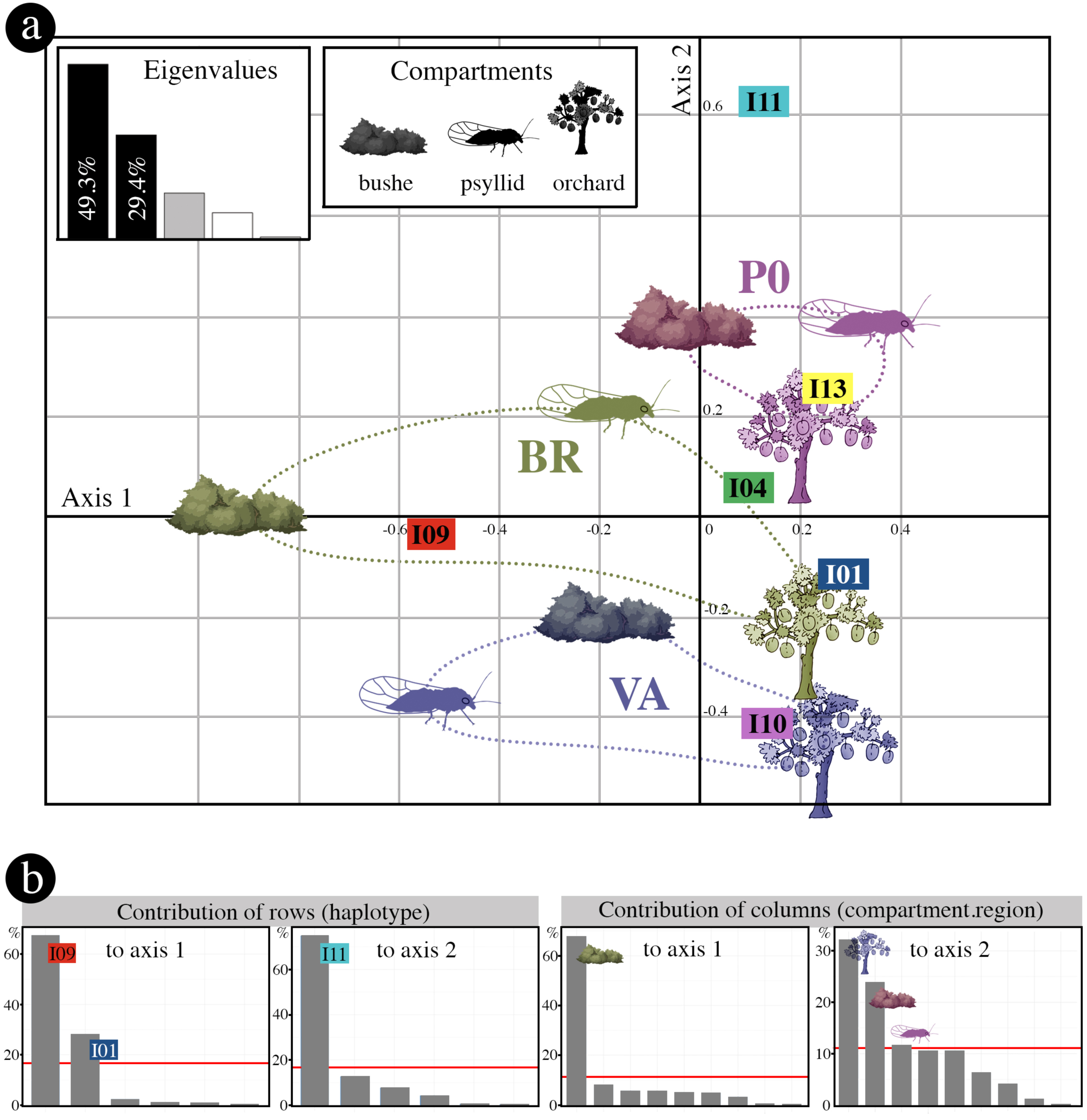
Correspondence analysis performed on the contingency table displaying the frequency distribution of the six major haplotypes in the three ecological compartments and the three regions. (a) Factorial map of the first two axes. Each compartment is represented by its symbol; the associated colours and ellipses are interpretation aids allowing to visualize the modalities associated with each region. The eigenvalue plot illustrates the percentage of variance explained by the different axes. (b) Contributions of rows and columns of the contingency table to the first two dimensions of the analysis. The red line represents the expected row or column contributions if the contributions were uniform. Any row or column above this threshold is a major contributor to the corresponding axis.

To assess the significance of these patterns, we first carried out multinomial logistic regressions, which evidenced a highly significant interaction (*p*=6.5×10^−7^; likelihood ratio test) between compartments and regions. Thus, we further studied these interactions by comparing the distributions of haplotypes between compartments for each region (Fig. 4a), and the distributions of haplotypes between regions for each compartment (Fig. 4b). In the PO region, there was no statistically significant difference (*p*=0.062; Fisher’s exact test) between the visually homogeneous compartment profiles with very close values of Simpson’s diversity index D (Fig. 4a). This region is characterized by a high prevalence of haplotype I11 in all three compartments, while this haplotype seems much rarer or absent in the compartments of the other two regions (Fig. 4a; Supplementary Tables S3 and S4). The BR region shows very contrasted haplotype frequency profiles among compartments (*p*<5×10^−6^; Fisher’s exact test) and strong differences between the values of the diversity index D. The high prevalence of haplotype I09 in the bushes, and conversely the high prevalence of haplotype I01 in the orchards contribute to this highly significant haplotype dissimilarity (Fig. 4a; Supplementary Tables S3 and S4). Within the VA region, the contrasted haplotypic composition (*p*≅6.3×10^−6^; Fisher’s exact test) is also reflected by the higher value of the diversity index among psyllids than among bushes or orchards. Within each compartment, the haplotypic composition differed significantly according to the growing region (Bushes: *p*<1×10^−8^; Psyllids: *p*≅5.3×10^−4^; Orchards; *p*<1×10^−8^) although the proportion of the main haplotypes was very similar across regions for the orchards (Fig. 4b) as mentioned above (Fig. 3b).

**Figure 4.**
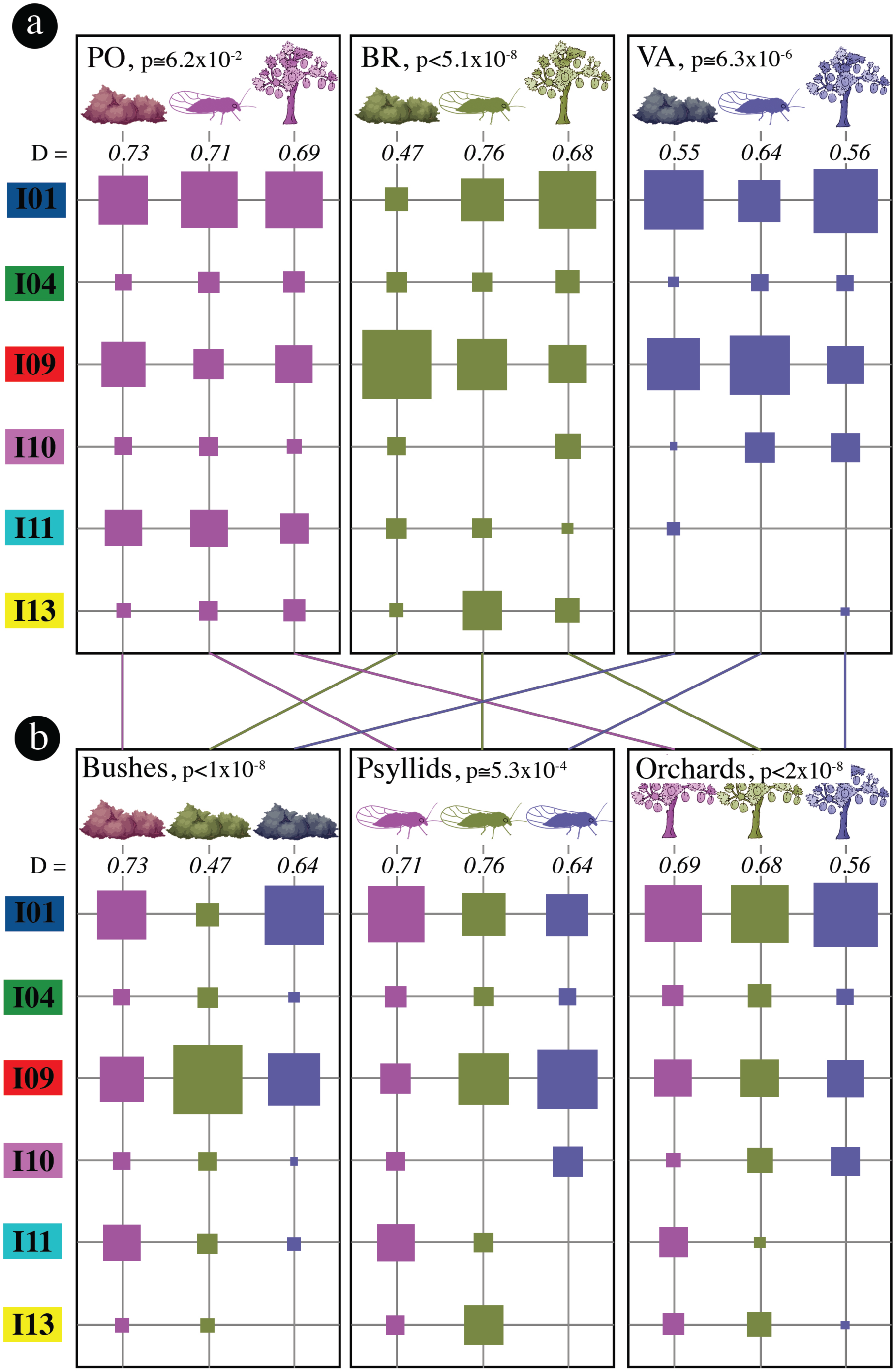
Relative proportions of the six major haplotypes in each of the three ecological compartments. The same data are either (a) grouped by region (PO: Pyrénées-Orientales, BR: Bas-Rhône, VA: Valence), or (b) grouped by ecological compartment. P-values correspond to Fisher’s exact test that haplotype profiles are identical either within a given region (in a), or within a given ecological compartment (in b). The values of Simpson’s diversity index D are italicised.

### Spatial patterns

In order to assess whether the sampled haplotypes were spatially structured within and between compartments, the relationship between genetic and geographical distances was tested at all distances, using a geostatistical method based on join counts^24^ and permutation tests. This approach is well adapted to an exploratory spatial analysis of epidemiological data because no assumption or prior knowledge of the processes of disease spread are needed (e.g., relative importance of several transmission pathways, distance and direction of spread, definition of population units)^39,40^. Because the abovementioned interaction between compartments and regions meant that the haplotype frequency distribution among compartments differed between the three growing regions, spatial analyses were performed independently within each region.

For the PO region (Figs. 5a and 6a), the tests indicated a strong spatial structure (i.e., a statistically highly significant excess of genetic similarity) up to 45-50 km for both the orchard and the bush samples, but not for the psyllid samples. The diversity indices D and D’ indicate that haplotypes are not genetically closer within than between compartments.

**Figure 5.**
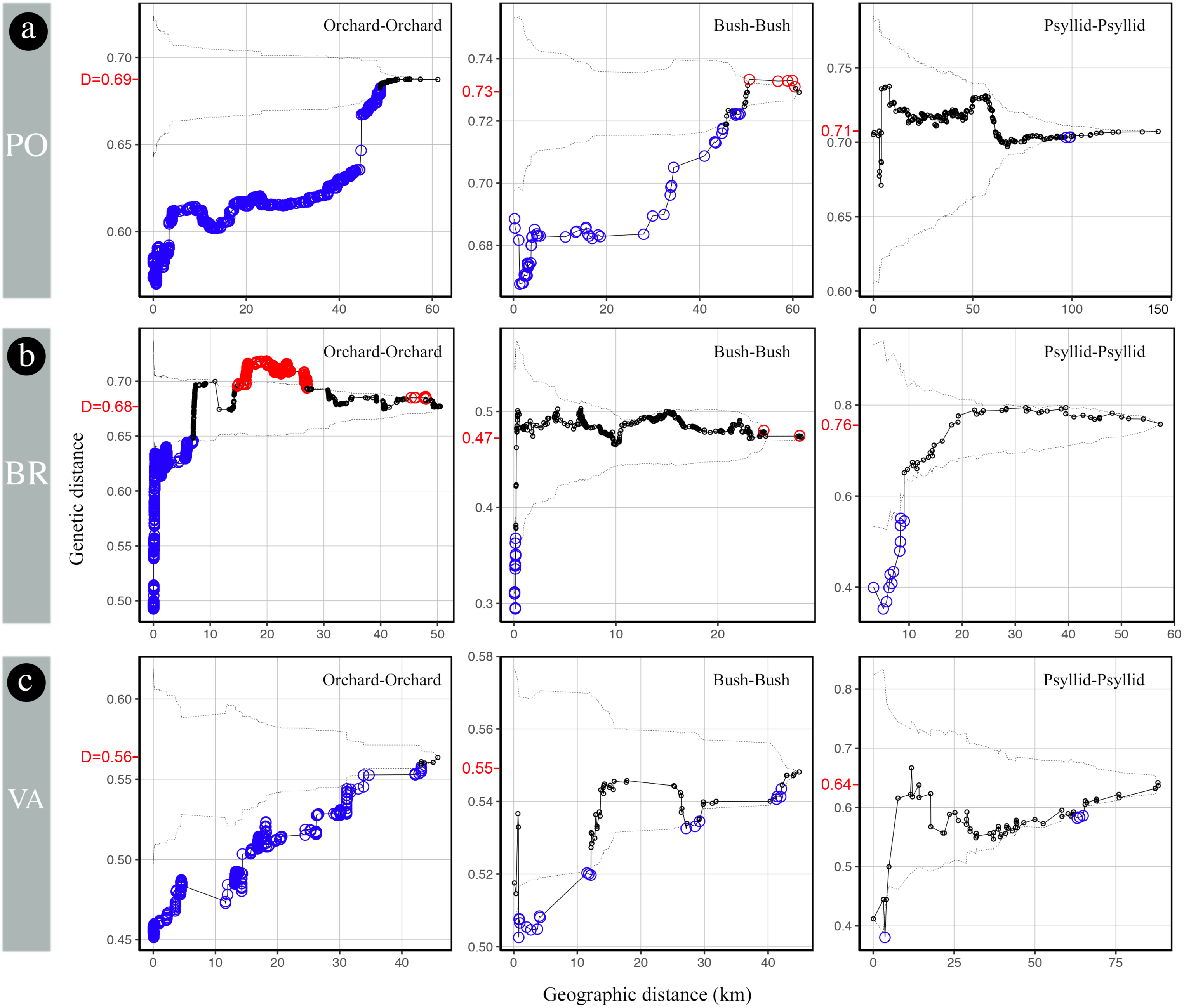
Mean pairwise genetic distances within increasing radii computed for all pairs of samples from each compartment in (a) Pyrénées-Orientales, (b) Bas-Rhône, and (c) Valence regions. Solid line, observed values; dashed lines, 95% confidence envelope under the null hypothesis of spatial independence. Bigger circles below (resp., above) the envelope indicate radii within which the mean pairwise genetic distance is significantly lower (resp., higher) than expected in the absence of geographical structure. The values of Simpson’s diversity index D are in red on the *y* axis.

**Figure 6.**
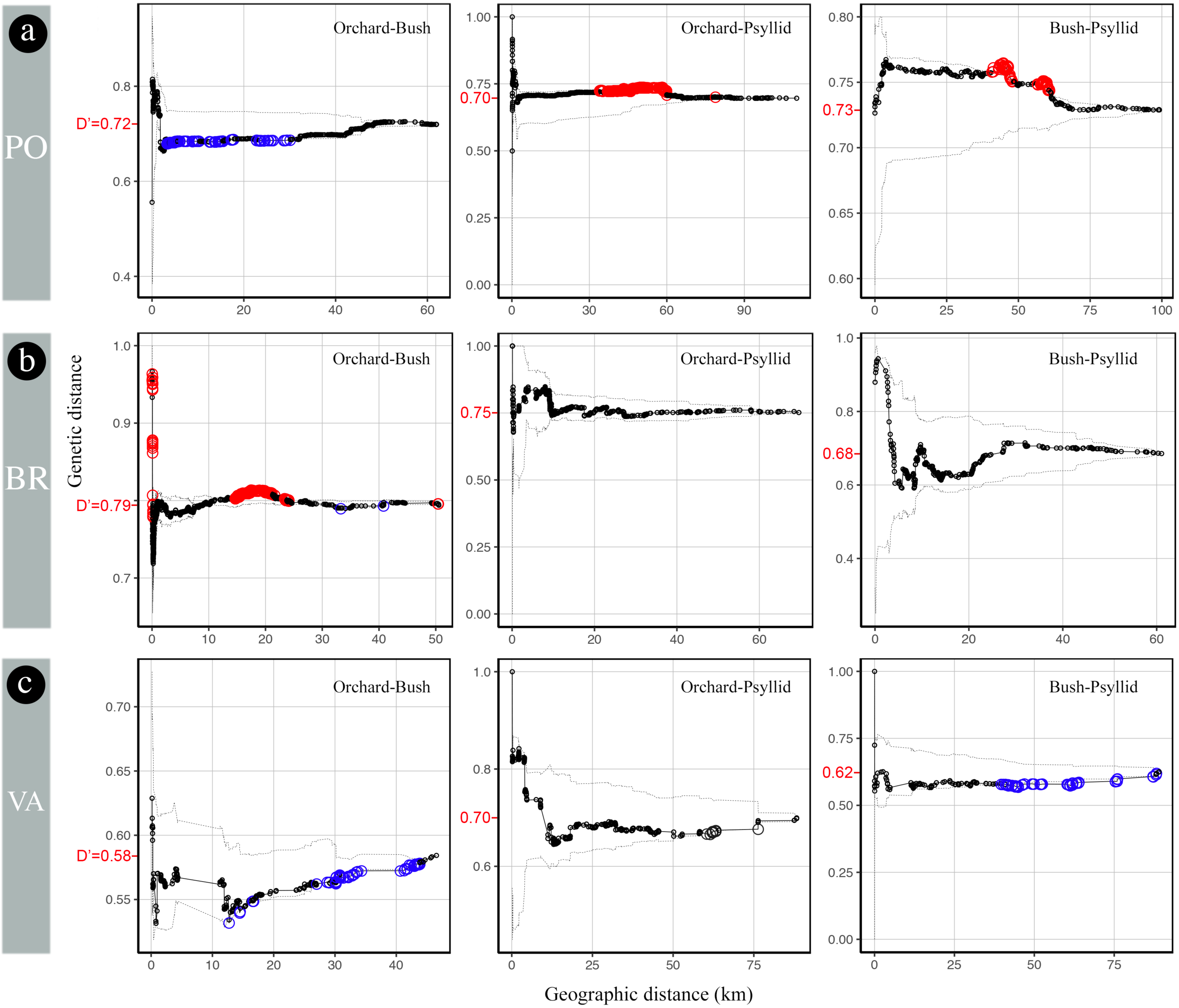
Mean pairwise genetic distances within increasing radii computed between all pairs of samples from different compartments in (a) Pyrénées-Orientales, (b) Bas-Rhône, and (c) Valence regions. Solid line, observed values; dashed lines, 95% confidence envelope under the null hypothesis of spatial independence. Bigger circles below (resp., above) the envelope indicate radii within which the mean pairwise genetic distance is significantly lower (resp., higher) than expected in the absence of spatial dependence between compartments. The marginally significant mid-distance genetic proximity between orchards and bushes in the PO region is an artefact of the permutation test caused by the highly significant genetic proximity of bush samples within 30 km (Fig. 5). The values of diversity index D’ are in red on the *y* axis.

For BR, the significant genetic proximity extended only up to 8 km for the orchard samples and 200 m for the bush samples, and the psyllid samples were genetically closer up to 9 km (Figs. 5b and 6b). The diversity indices D’ for the three pairs of compartments varied in the range 0.68-0.79. This latter value for all orchard-bush pairs of samples is higher than the diversity indexes D within these two compartments (0.47 and 0.68), confirming the lack of genetic proximity between orchards and bushes.

For VA, the significant genetic proximity extended up to 35 km for the orchards, only up to 12 km for the bush samples, and the psyllid samples were genetically closer within 4 km (Figs. 5c and 6c). The diversity indices D’ for the three pairs of compartments varied in the range 0.58-0.70. This latter value for all orchard-psyllid pairs of samples is higher than the diversity indices D within these two compartments (0.56 and 0.64), confirming the overall genetic differentiation between samples from orchards and psyllids.

A striking common feature across all three regions is the absence of significant genetic proximity between different nearby compartments (Fig. 6a), which suggests limited spread at short spatiotemporal scales.

## DISCUSSION

In this work, we used complementary statistical approaches combining genetic data and geographical distances to identify spatial genetic patterns that provide insights into various scenarios regarding the eco-epidemiology of ‘*Ca*. P. prunorum’, including the role of humans and vectors and the associated scales of pathogen spread.

Several results of our study strengthen previous information on the functioning of the pathosystem. Firstly, we show that the pathogen is highly prevalent in all of the 61 tested bushes, confirming previous reports from other European countries^32,34^. Secondly, our study provides good estimates of the phytoplasma prevalence among French populations of immigrant psyllids (3.2% -6.4%; Table 1). Overall, this low infection rate is in line with those estimated in previous studies with psyllids captured also in the natural environment in spring in France, Germany, Austria, and Bulgaria^33^. Much higher infection rates in psyllids were reported locally in Italy, Switzerland, and Spain, but they might be caused by specific environmental conditions (e.g., very high level of psyllid populations in blackthorn and in untreated orchards in Swiss Valais; higher host attractiveness or palatability, as seen for Japanese plum trees in Italy), or specific local psyllid populations as seen with vectors of ‘*Ca*. P. mali’^33^. Because phytoplasma prevalence is much lower in psyllids than in bushes at the regional scale, phytoplasma acquisition from blackthorn appears to be inefficient.

Our study also brings new information on the vectors of ‘*Ca*. P. prunorum’. The discovery that *C. pruni* is actually a complex of two cryptic species^37,38^ raised questions on their respective distribution area and vector competence. In the present study, we observed balanced species proportions (Supplementary Table S5) in the PO region (59% A) and in the VA region (55% A). In contrast, we found almost only species A in the BR region (97% A), and species B was the only species described in northern Europe^33,34,41^. The PO and VA regions may thus be part of a contact zone between the two species. Given the apparent absence of species A in areas where ‘*Ca*. P. prunorum’ is found in wild and cultivated *Prunus*^33,34^, there is little doubt that species B is a vector of this pathogen; however, the vectorial competence of species A had not been previously documented. The present study strongly supports that species A is also a vector of ‘*Ca*. P. prunorum’: the six major haplotypes are present in both psyllid species, pathogen prevalence is similar in psyllids of species A and B and, more importantly, in bushes from the three regions (including BR where species B is very rare). Dedicated transmission trials with *C. pruni* species A would put this hypothesis to the test.

No previous study addressed the key question of the spatial scale of ESFY epidemics. Here, we show unambiguously a sub-regional structure of the pathogen haplotypes, and provide new insights into the main drivers of the epidemic. In particular, we demonstrate that the haplotype distribution of the pathogen in the bushes differs very significantly between the three regions (Fig. 4b), which means that there are few or no natural connections (i.e., psyllid dispersal) between the three major French regions growing stone fruit trees. One of the crucial unknowns in the different scenarios illustrated in Fig. 1 and Supplementary Fig. S1 was the scale of vector dispersal between pathogen acquisition and transmission. Our study shows that pathogen haplotypes are more similar than expected (i.e., sites are more epidemiologically connected) within a radius of at most 50 km (Fig. 5), which explains the observed strong genetic structure between the regions, which are >100 km apart. The most important result is arguably the absence of spatial dependence between the haplotypes sampled from bushes and orchards within each of the three regions despite (i) the strong spatial structure of the haplotypes within each compartment, and (ii) the high statistical power to detect this dependence. This means that the main drivers of the epidemic in the studied regions are neither the direct transmission of pathogens between bushes and orchards by psyllids (scenario 2), nor the transmission by infectious vectors which would successively land in bushes and then in surrounding orchards (scenario 6). In the same vein, this means that scenario 14 (i.e., transmission per bounces of infectious psyllids from orchards to nearby blackthorns, see Supplementary Figure 1) is also rejected. However, this result is unexpected because this scenario seemed compatible with the biology of the insect (i.e., insects flying in early spring, landing randomly and passing from one plant to another to find their partners and breeding sites). The absence of spatial dependence between the haplotypes sampled from bushes and orchards also excludes a local or philopatric version of scenario 4, in which bushes and neighbouring orchards would be connected through shelter sites. The presence of all major haplotypes in all compartments (Fig. 4b) and the similarities in haplotype distributions between compartments at the regional scale (Figs. 3 and 4a)—in particular for PO where more samples were collected (Table 1)—implies regional gene flows between wild and cultivated *Prunus*, i.e. scenario 4 is necessary to explain this observed pattern. However, this scenario is not sufficient to explain that the haplotype composition of ‘*Ca*. P. prunorum’ is more similar across regions for orchards than for the wild compartments (Figs. 3 and 4). This observation implies a significant contribution of inter-regional exchanges of infected plants (for planting) to the long-distance spread of the disease in orchards, i.e. scenario 8 is also necessary to explain this observed pattern. A logical consequence is that a more local version of this scenario (i.e., scenario 7_a+b_) must also contribute to the long spatial range of the haplotype similarity between orchards within each region (Fig. 5). The observed patterns are not directly informative on scenarios 1, 3 and 5. However, both scenarios 1 and 3 are unlikely to contribute significantly to the observed patterns because they rely on pathogen acquisition in orchards. Now, except in organic farming, psyllids are very rarely observed in orchards because of probable combined effect of insecticides and lower attractiveness of cultivated *Prunus* compared to wild *Prunus*). Futhermore, the stone fruit trees are much less abundant in the natural habitat than wild *Prunus*, and much less infected, as shown in our study. All these factors contribute to making this cultivated compartment a weak reservoir of inoculum and/or insects. Finally, successive transmission events within orchards by infectious (mature) adults (scenario 5) is compatible with the persistent transmission mode. To summarise, this field study supports and refines previous hypotheses derived from laboratory experiments^9^ by showing (i) that exchanges of infected plants contribute to disease spread between regions (a risk pointed out by several works^30,41^, and (ii) that pathogen spread within the region involves acquisition (mainly from wild *Prunus*) by larvae and immature adults, a first migration before the summer to one of the regional shelter hubs (conifers in altitude within a few tens of kilometres), a second migration after the winter and (possibly multiple) transmissions within *Prunus* orchards after an effective latency of 8 months. Therefore, ESFY may be seen as a self-sustaining natural pathosystem that accidentally impacts stone fruit species, in particular in apricot orchards where ESFY is essentially a monocyclic disease (i.e. with no secondary spread within the cultivated compartment). Such an epidemic cycle is an extremely unusual feature for plant diseases, and evoke vector-borne zoonoses like Lyme disease or West Nile fever ^42^.

The new insights on the pathosystem fill some gaps identified in an EFSA pest risk assessment^43^, such as the origin of contaminations in European orchards. Indeed, this work highlights the prominent role of wild *Prunus* as a source of infection for apricot trees. Given the endemic nature of this disease in Europe in the wild habitat^32^, and the very wide distribution area of the psyllid vectors^44^, there is little doubt that the disease cycle is similar in most diseased areas. Nevertheless, our study does not rule out the possibility that attractive *Prunus* orchards could be a significant source of inoculum in some particular situations (e.g., abandoned or organic orchards, heavily infested by psyllids and the pathogen). In the current state of knowledge of the pathosystem, once established in the natural environment, it would be unrealistic to consider eradicating the disease at any scale (farm, production basin, country). Disease control should thus focus on the protection of orchards against the psyllid vectors during the key period of their return flight after overwintering, and on the production of healthy plants for planting (in particular by the protection of nurseries).

The approach used in this work could be applied to other pathosystems, in particular in the case of plant diseases due to vector-borne bacteria (e.g., ‘*Candidatus* Phytoplasma spp.’, ‘*Candidatus* Liberibacter spp.’, *Xylella fastidiosa*), for which disease management strategies would strongly benefit from insights into the epidemiological role of wild plants or the scale of disease dispersal^6,8,10,33,45,46^. The top-down exploratory approach that we applied here can be implemented relatively easily, using a single molecular marker for a clonal organism sampled from wild and cultivated compartments and a robust multiscale statistical approach involving correspondence and join-count analyses. Further insights may be gained on the complex interaction network among host plants and insect vectors by the use of landscape genetic approaches^23,47,48^ to analyse the haplotypes of ‘*Ca*. P. prunorum’ from insects collected in, and at various distances from, shelter sites.

## METHODS

### Study area and sampling

In each growing area, we sampled mainly (90%) apricot (*Prunus armeniaca* L.) orchards (Supplementary Table S1); the remaining 10% of sampled orchards consisted of cultivated myrobalan plum (*Prunus cerasifera* Ehrh.), European Plum (*Prunus domestica* L.), Japanese plum (*Prunus salicina* Lindl.), and peach (*Prunus persica* (L.) Batsch) trees. Most of the samples were collected on symptomatic trees during autumns and winters 2010 and 2011 (Supplementary Table S1). We also included samples obtained in previous years (2007, 2008, and 2009). From each tree, we sampled 2-3 lignified shoots from different main branches. After molecular tests to assess the presence of the phytoplasma (see below), four to six plots were chosen in each region for a more comprehensive sampling (Supplementary Table S1). This was done from winter 2010 to early spring 2011. In this second sampling, all the symptomatic trees were collected (thus, some trees were sampled several times for confirmation of previous molecular analyses; see below); to estimate the number of asymptomatic trees (data unknown at the beginning of the study, but crucial to estimate the potential role of orchards as inoculum reservoir), one every three trees was also collected (i.e., systematic sampling). Between 78 and 244 trees per plot were analysed, depending on the size of the plots (Supplementary Table S1). A total of 2,656 samples collected in 69 different orchards were used in the study (Supplementary Table S1). At the same time, we sampled wild bushes of blackthorn (*Prunus spinosa* L.) or myrobalan around the plots, at up to a few dozen kilometres (Supplementary Figs. S2, S3 and S4). We collected on average 21 basal branches in 61 bushes, for a total of 1,114 samples (Tables 1; Supplementary Table S1). We also collected as many mature *C. pruni* as possible using a beating tray (80×80 cm), conserving them immediately in 96% ethanol until DNA extraction. Phytoplasma-carrying insects collected several years before in the three regions were also included in the analysis (Supplementary Table S1). Thus, a total of 2,572 psyllids sampled from 71 different bushes were analysed (Tables 1; Supplementary Table S1). We recorded the GPS coordinates of all collected samples, except for the systematically sampled orchards where we attributed a unique GPS coordinate—corresponding to the centre of each plot—to all the corresponding samples (Supplementary Figs. S2, S3 and S4).

### Genetic analyses

The protocol used for the total DNA extraction from plant samples was adapted from Ahrens and Seemüller^49^. Briefly, for each plant sample, the phloem was isolated by removing the outer bark with a knife and by scraping off the layer of vascular tissue with a scalpel. Phloem tissue was then ground in individual bags (0.5 g per bag). All the *Prunus* samples were analysed individually. Then, DNA from plant samples was purified using the CTAB method^50^ in 1.5 ml tubes. DNA pellets were diluted in 100 µl of pure water. Total DNA of individual psyllids was purified from whole bodies and each psyllid DNA sample was assigned to species A or B by amplifying the Internal Transcribed Spacer 2 (ITS2) as previously^37^. No individual showed two bands, demonstrating that the sample was devoid of hybrids or contamination between species.

To select samples for sequencing, ‘*Ca*. P. prunorum’ was detected in the insect and plant samples by using the ESFYf/r primers in a specific and sensitive PCR-based method as described in Yvon et al.^51^. Then, we tried to sequence the 1,328 positive samples at the immunodominant membrane protein (*imp*) gene locus, which was shown to be highly variable for ‘*Ca*. P. prunorum’^38^ and assumed to be present at a single copy per genome based on the known genome sequence of ‘*Ca*. P. mali’^52^. DNA amplification performed well for almost all psyllid samples, but failed for more than a quarter of the plant samples (Table 1), which we attributed to the presence of putative inhibitors like polyphenols^53^ in the unevenly infected woody material. Successfully amplified *imp* DNA was purified and Sanger-sequenced in both directions by Genewiz (Takeley, UK). Chromatograms were trimmed, assembled, and aligned using the Muscle algorithm, and visually checked under Geneious version 5.5 (http://www.geneious.com). Sequences were deposited in GenBank (accession n° MN116709 to MN116718; Supplementary Table S6). Single nucleotide polymorphisms (SNP) between individual sequences were detected, and ‘*Ca*. P. prunorum’ haplotypes were defined as in Danet et al.^35^. When previously undescribed SNPs were detected, we performed a second independent extraction from the same sample followed by amplification and sequencing to ascertain the new *imp* sequence. To represent genealogical relationships among sequences, we used the POPART software^54^ (v.1.7) to build a haplotype network using the integer neighbour-joining (IntNJ) algorithm, which is well adapted for low-divergence data sets. Each infected individual contained a single *imp* haplotype, except for six cultivated trees in which two different haplotypes were found (either from different branches or in different years) and kept for the analysis.

### Statistical analysis

All statistical analyses were performed using R 3.4.0^55^. A correspondence analysis was performed on the contingency table (Supplementary Table S3) using the function *coa* in the package *ade4*^56^, and we visualized the results with the package *factoextra*^57^. The contingency table was also directly visualized using the function *table*.*value* in *ade4* to uncover specific association patterns. To test whether the distribution of haplotypes among compartments differed between the three regions, we carried out multinomial regressions with the package *nnet*, and we tested the interaction between compartments and regions by comparing (using a likelihood ratio test) the complete model (including the main effects and their interaction) with the model without the interaction. Fisher’s exact tests with simulated p-value (based on 10^8^ replicates) were used to test the homogeneity of the distributions of haplotypes between compartments (within each region), and of haplotypes between regions (within each compartment).

In the geostatistical analyses, the genetic distance between two samples was fixed at 0 if the samples had the same haplotype and at 1 if their haplotypes differed. This definition corresponds to the join count between elements of different classes of nominal data^24^. In cases where two different haplotypes were detected in the same tree, their geographical distance was set at 0 (and their genetic distance at 1). In the case of a single compartment with *n* samples, the geographical distances between the *n*(*n –* 1)/2 pairs of samples were first calculated. Then, for each calculated distance *d*, we computed the average genetic distance *D*_*d*_ between the *k*_*d*_ pairs of samples separated by a geographical distance less than or equal to *d* (i.e., within a radius *d*). The average genetic distance was defined as:

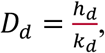

with *h*_*d*_ the number of pairs of samples with different haplotypes among the *k*_*d*_ pairs. The confidence intervals (here, at level *α* = 0.05) were obtained from *N* (here, *N* = 10,000) random permutations of the haplotypes of the *n* samples. We calculated the *N* average genetic distances within each distance *d*, and the lower (respectively, upper) limit was defined as the genetic distance with rank N*α*/2 (resp., *N*(1 – *α*/2)) among the *N* random genetic distances. A significant reduction in genetic diversity at the beginning of the curve (i.e., for the smaller radii) is expected when haplotypes are spatially clustered, and the distance at which this reduced genetic diversity becomes non-significant indicates the spatial extent of genetic similarity among samples. Significant genetic distances are interpreted as false positives when nonsignificant genetic distances are observed at shorter geographical distances (except when these shorter distances are associated with a very low statistical power, i.e., wide confidence envelopes). Considering the cumulative number of pairs separated by a distance less than or equal to *d*, rather than the number of pairs falling in distance classes defined by intervals, provides more powerful tests and more stable curves.

When *d* is equal to the maximum distance *d*_*max*_ between two samples, we have:

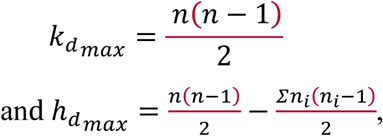

where *n*_*i*_ is the number of samples with haplotype *i* and the summation is over all the haplotypes. Thus, the value of 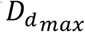 is:

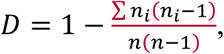

which is Simpson’s diversity index^61^.*D* = 0 if all the samples have the same haplotype, and *D* = 1 if all the samples have different haplotypes. As 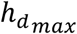 does not change when the haplotypes are permuted, the lower and upper limits of the confidence interval are equal to *D* when *d = d*_*max*_. When *d* < *d*_*max*_, *D*_*d*_ can be considered as Simpson’s diversity index restricted to the pairs of samples separated by a geographical distance less than or equal to *d*.

In the case of two different compartments with *n*_1_ and *n*_2_ samples, a similar procedure was applied to the *n*_1_*n*_2_ pairs consisting of one sample of each compartment with the statistics

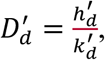

where 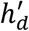 and 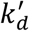 have the same definition as *h*_*d*_ and *k*_*d*_ except that the two samples belong to two different compartments. However, for the computation of the confidence intervals, only the haplotypes of the less structured compartment were randomly permuted to prevent false positive tests caused only by breaking the structure of the most structured compartment by permutation^58^. When the less structured compartment was also significantly structured, the tests were interpreted conservatively, i.e., interpreting as not statistically significant the points bordering the limits of the confidence envelope. When *d = d*_*max*_, the value of 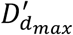 is:

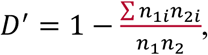

where *n*_1*i*_ and *n*_2*i*_ are the numbers of samples with haplotype *i* in each of the two groups. *D′ = 0* if the two groups have the same unique haplotype and *D′ = 1* if the two groups have no common haplotype.

## Data availability

## Acknowledgements

We are very grateful to J. Peyre for her valuable technical assistance, and the following individuals for their contributions to the collection of psyllid or plant samples: N. Courtieu & J.-M. Broquaire (SICA Centrex), N. Galabert & J. Delnatte (SICA L’Edelweiss), B. Rouillé (SRPV-PACA), E. Navarro (Terroir de Crau), P. Delon (CA-Gard), E. Falezan (GIE-Tain l’Hermitage), P. Exbrayat (CA-Drôme), M. Léon-Chapoux & V. Delaunay (SEFRA), and the fifteen growers involved in this study. Part of this work benefited from a postdoctoral grant to JP funded by INRA-CIRAD grant SDIPS (Speciation and molecular Diagnosis of Insect Pest Species complexes). Field and molecular work for this study were supported by the project Prima phacie funded by EFSA grant agreement CFP/EFSA/PLH/2009/01.

## Author Contributions

Conceived and designed the experiments: FB, GT, JP, GL, NS; performed the experiments: VMJ, JP, GL, NS; analysed the data: VMJ, FB, GT, NS; contributed reagents/materials/analysis tools: VMJ, FB, JP, NS; wrote the paper: VMJ, FB, GT, JP, GL, NS.

## Supplementary Information

**S1 Figure**. Complete sets of eco-epidemiological scenarios illustrating the pathways through which ‘*Ca*. P. prunorum’ might be spread in orchards and wild habitat. Bi: bushes (i.e. wild *Prunus* = host plants); Ci: conifers (i.e. shelter plants); Ni: nurseries; Oi: orchards. In red: infected plants or infectious psyllids; in green: non-infected plants. Contamination of cultivated trees at more or less great spatio-temporal scales can be imagined according to the scenarios: contamination of an apricot tree by a psyllid from a nearby tree (S1) or a nearby bush (S2); contamination by an immigrant who has acquired phytoplasma on an infested tree (S3) or in a bush (S4) the previous year; multiple contaminations by the same infectious psyllid of trees from the same orchard (S5), or from a bush and then a tree (or trees) from a nearby orchard (S6); contamination of orchards by plants from nearby nurseries (S7a, S7b); contaminations of different orchards by plants from the same nursery, not necessarily of the same geographic area (S8). The contamination of wild *Prunus* can be illustrated by scenarios identical to those of apricot trees (scenarios S9 to S14).

**S2 Figure**. Maps of the spatial distribution of samples for each compartment (bush, psyllid, and orchard) surveyed in the three growing regions of the study. n: number of *imp* sequences successfully genotyped in each compartment.

**S3 Figure**. Map of the spatial distribution of samples from each compartment (bush, psyllid, and orchard) surveyed in the Pyrénées-Orientales (PO) region. n: number of *imp* sequences successfully genotyped in each compartment. A small random noise was added to the sample coordinates to avoid overlapping.

**S4 Figure**. Detailed map of the spatial distribution of samples from each compartment (bush, psyllid, and orchard) surveyed in the Bas-Rhône (BR) region. n: number of *imp* sequences successfully genotyped in each compartment. A small random noise was added to the sample coordinates to avoid overlapping.

**S5 Figure**. Detailed map of the spatial distribution of samples from each compartment (bush, psyllid, and orchard) surveyed in the Valence (VA) region. n: number of *imp* sequences successfully genotyped in each compartment. A small random noise was added to the sample coordinates to avoid overlapping.

**S1 Table**. Summarized data for all samples surveyed in the study, ordered by region (PO, BR, VA) and ecological compartment (bush, psyllid, orchard). For each compartment in each region, statistics (total number, minimum/maximum/mean values) are given for the variables: N, total number of samples collected in each region; #ESFY+, total number of samples found positive by PCR; %ESFY+, mean percentage of samples found positive by PCR; #IMP, number of samples successfully genotyped for the *imp* gene. A more concise summary of these data is provided in Table 1.

**S2 Table**. Contingency table of the frequency distribution of all haplotypes successfully genotyped in each ecological compartment (bush, psyllid, and orchard) and each of the three growing regions. Haplotypes never described in previous studies are italicized. The legend in Fig. 2 explains how these new haplotypes were named. Asterisks indicate changes in the amino acid sequence.

**S3 Table**. Contingency table displaying the frequency distribution of the six major *imp* haplotypes in each ecological compartment (bush, psyllid, and orchard) and each of the three growing regions of the study. PO: Pyrénées-Orientales; BR: Bas-Rhône; VA: Valence. Very rare haplotypes (i.e., with frequency <1% combining all regions) were excluded from the analyses. See Supplementary Table S2 for more details.

**S4 Table**. Relative proportions of the six major haplotypes in the three ecological compartments and the three regions. PO: Pyrénées-Orientales, BR: Bas-Rhône, VA: Valence. This table is summarized in Fig. 4.

**S5 Table**. Number of major *imp* haplotypes genotyped for each species of the *Cacopsylla pruni* complex, all regions combined, and region by region. PO: Pyrénées-Orientales; BR: Bas-Rhône; VA: Valence. #A/B: total number of psyllids collected from species A and/or B. #ESFY+: total number of samples found positive by PCR. %A/B: proportion of psyllids of each psyllid species. %ESFY+: proportion of each psyllid species among the PCR-positive psyllids.

**S6 Table**. List of the *imp* haplotypes described to date, with their GenBank accession numbers.

