## Supplemental Information for "Multi-scale spatial genetic structure of a vector-borne plant pathogen in orchards and wild habitat"

**Supplementary Information**

**S1 Figure.** Complete sets of eco-epidemiological scenarios illustrating the pathways through which '*Ca*. P. prunorum' might be spread in orchards and wild habitat. Bi: bushes (i.e. wild *Prunus* = host plants); Ci: conifers (i.e. shelter plants); Ni: nurseries; Oi: orchards. In red: infected plants or infectious psyllids; in green: non-infected plants. Contamination of cultivated trees at more or less great spatio-temporal scales can be imagined according to the scenarios: contamination of an apricot tree by a psyllid from a nearby tree (S1) or a nearby bush (S2); contamination by an immigrant who has acquired phytoplasma on an infested tree (S3) or in a bush (S4) the previous year; multiple contaminations by the same infectious psyllid of trees from the same orchard (S5), or from a bush and then a tree (or trees) from a nearby orchard (S6); contamination of orchards by plants from nearby nurseries (S7a, S7b); contaminations of different orchards by plants from the same nursery, not necessarily of the same geographic area (S8). The contamination of wild *Prunus* can be illustrated by scenarios identical to those of apricot trees (scenarios S9 to S14).

**S2 Figure.** Maps of the spatial distribution of samples for each compartment (bush, psyllid, and orchard) surveyed in the three growing regions of the study. n: number of *imp* sequences successfully genotyped in each compartment.

**S3 Figure.** Map of the spatial distribution of samples from each compartment (bush, psyllid, and orchard) surveyed in the Pyrénées-Orientales (PO) region. n: number of *imp* sequences successfully genotyped in each compartment. A small random noise was added to the sample coordinates to avoid overlapping.

**S4 Figure.** Detailed map of the spatial distribution of samples from each compartment (bush, psyllid, and orchard) surveyed in the Bas-Rhône (BR) region. n: number of *imp* sequences successfully genotyped in each compartment. A small random noise was added to the sample coordinates to avoid overlapping.

**S5 Figure.** Detailed map of the spatial distribution of samples from each compartment (bush, psyllid, and orchard) surveyed in the Valence (VA) region. n: number of *imp* sequences successfully genotyped in each compartment. A small random noise was added to the sample coordinates to avoid overlapping.

**S1 Table.** Summarized data for all samples surveyed in the study, ordered by region (PO, BR, VA) and ecological compartment (bush, psyllid, orchard). For each compartment in each region, statistics (total number, minimum/maximum/mean values) are given for the variables: N, total number of samples collected in each region; #ESFY+, total number of samples found positive by PCR; %ESFY+, mean percentage of samples found positive by PCR; #IMP, number of samples successfully genotyped for the *imp* gene. A more concise summary of these data is provided in Table 1.

**S2 Table.** Contingency table of the frequency distribution of all haplotypes successfully genotyped in each ecological compartment (bush, psyllid, and orchard) and each of the three growing regions. Haplotypes never described in previous studies are italicized. The legend in Fig. 2 explains how these new haplotypes were named. Asterisks indicate changes in the amino acid sequence.

**S3 Table.** Contingency table displaying the frequency distribution of the six major *imp* haplotypes in each ecological compartment (bush, psyllid, and orchard) and each of the three growing regions of the study. PO: Pyrénées-Orientales; BR: Bas-Rhône; VA: Valence. Very rare haplotypes (i.e., with frequency <1% combining all regions) were excluded from the analyses. See Supplementary Table S2 for more details.

**S4 Table.** Relative proportions of the six major haplotypes in the three ecological compartments and the three regions. PO: Pyrénées-Orientales, BR: Bas-Rhône, VA: Valence. This table is summarized in Fig. 4.

**S5 Table.** Number of major *imp* haplotypes genotyped for each species of the *Cacopsylla pruni* complex, all regions combined, and region by region. PO: Pyrénées-Orientales; BR: Bas-Rhône; VA: Valence. #A/B: total number of psyllids collected from species A and/or B. #ESFY+: total number of samples found positive by PCR. %A/B: proportion of psyllids of each psyllid species. %ESFY+: proportion of each psyllid species among the PCR-positive psyllids.

**S6 Table**. List of the *imp* haplotypes described to date, with their GenBank accession numbers.

**S1 Figure.** Complete sets of eco-epidemiological scenarios illustrating the pathways through which '*Ca.* P. prunorum' might be spread in orchards and wild habitat

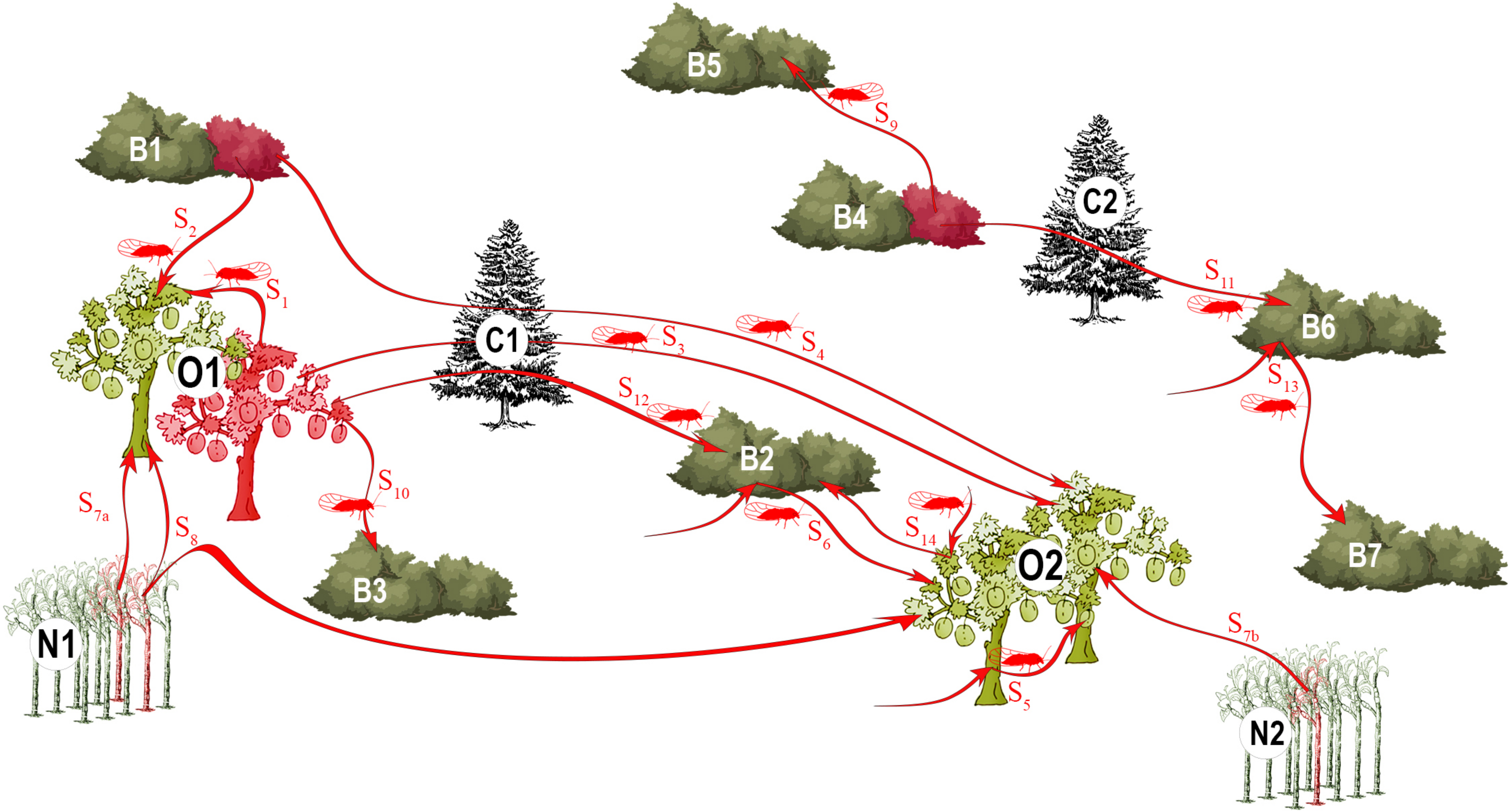

**Figure S2.** Maps of the spatial distribution of samples for each compartment (bush, psyllid, and orchard) surveyed in the three growing regions of the study.

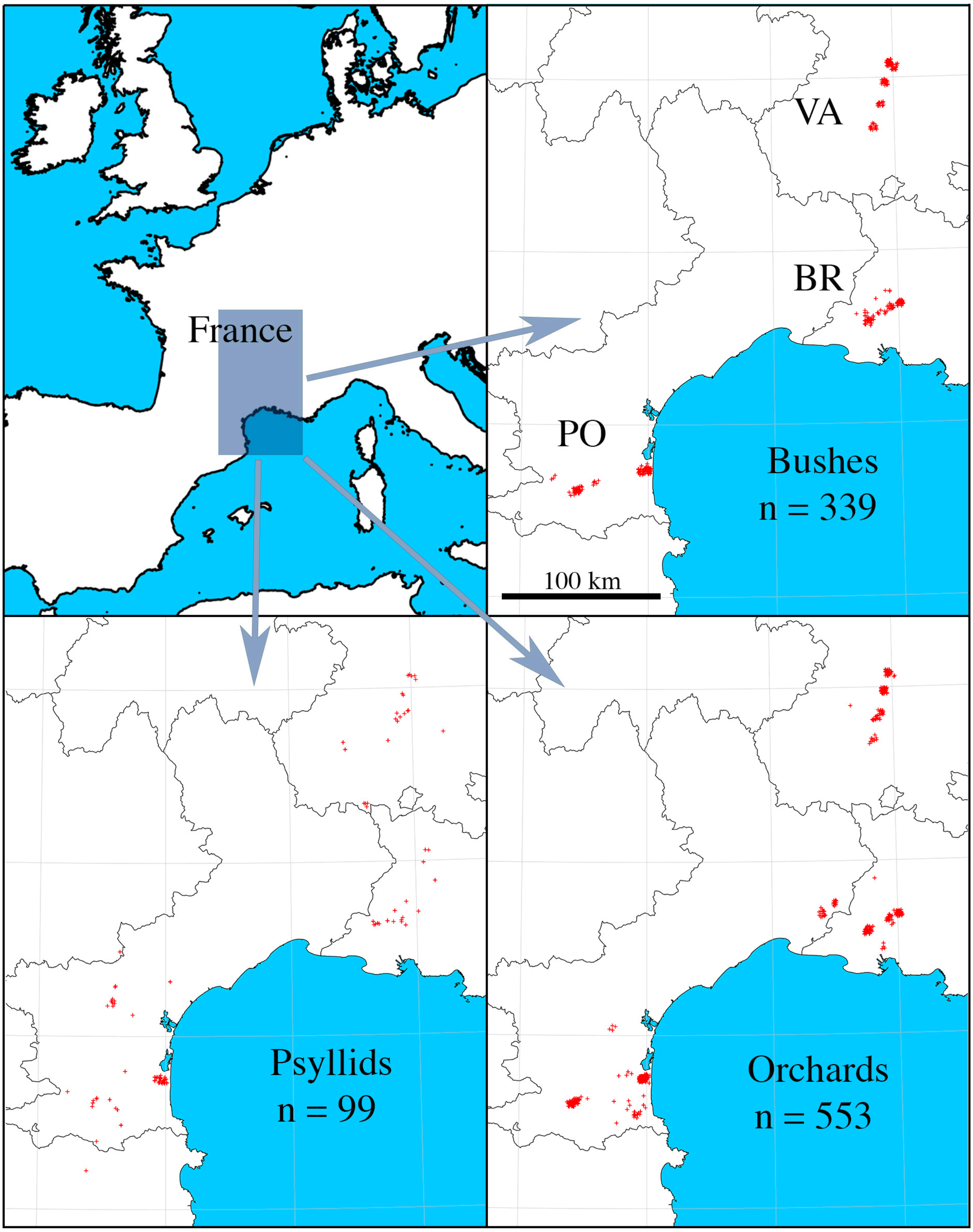

**Figure S3.** Detailed map of the spatial distribution of samples from each compartment (bush, psyllid, and orchard) surveyed in the Pyrénées-Orientales (PO) region.

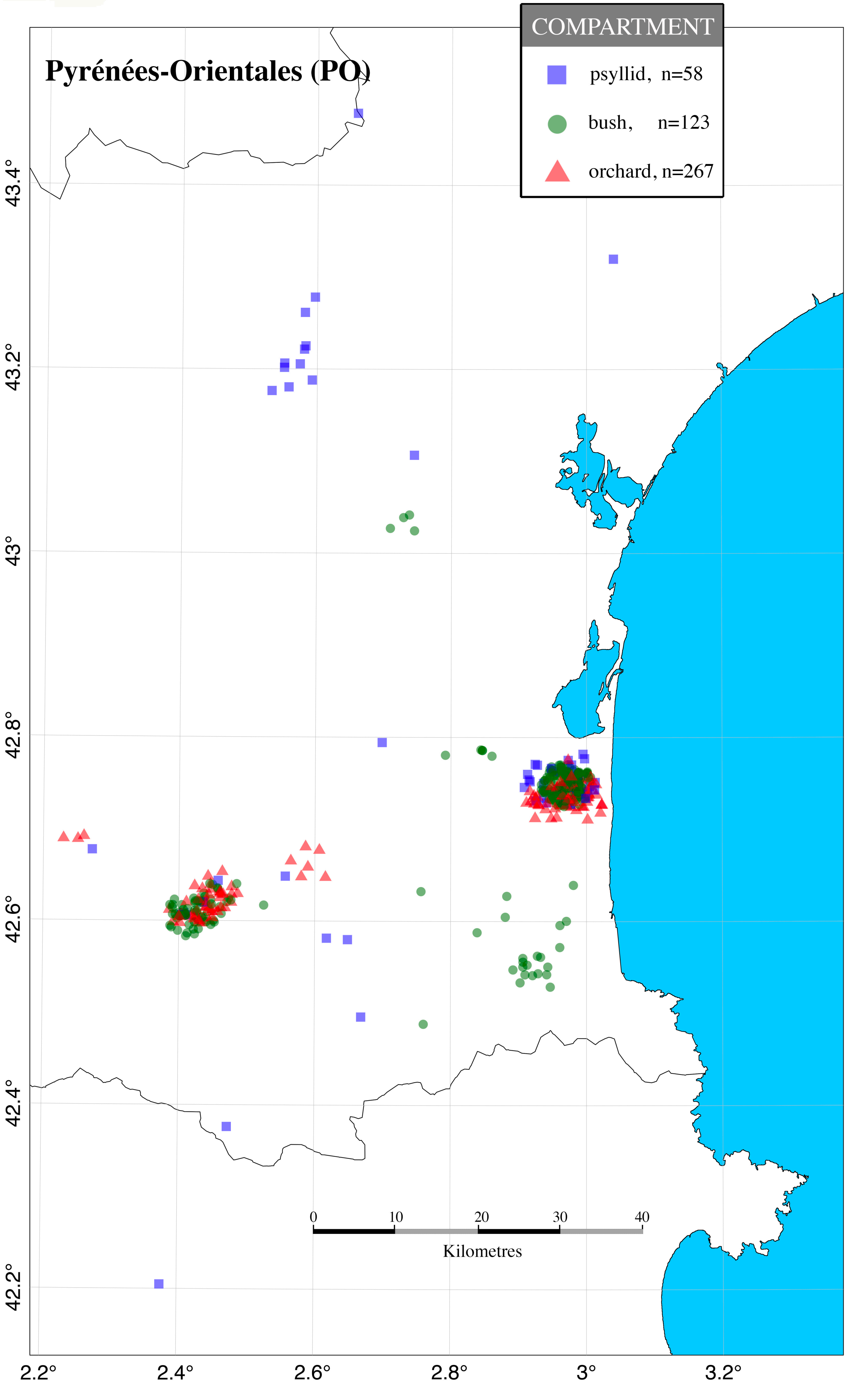

**Figure S4.** Detailed map of the spatial distribution of samples from each compartment (bush, psyllid, and orchard) surveyed in the Bas-Rhône (BR) region.

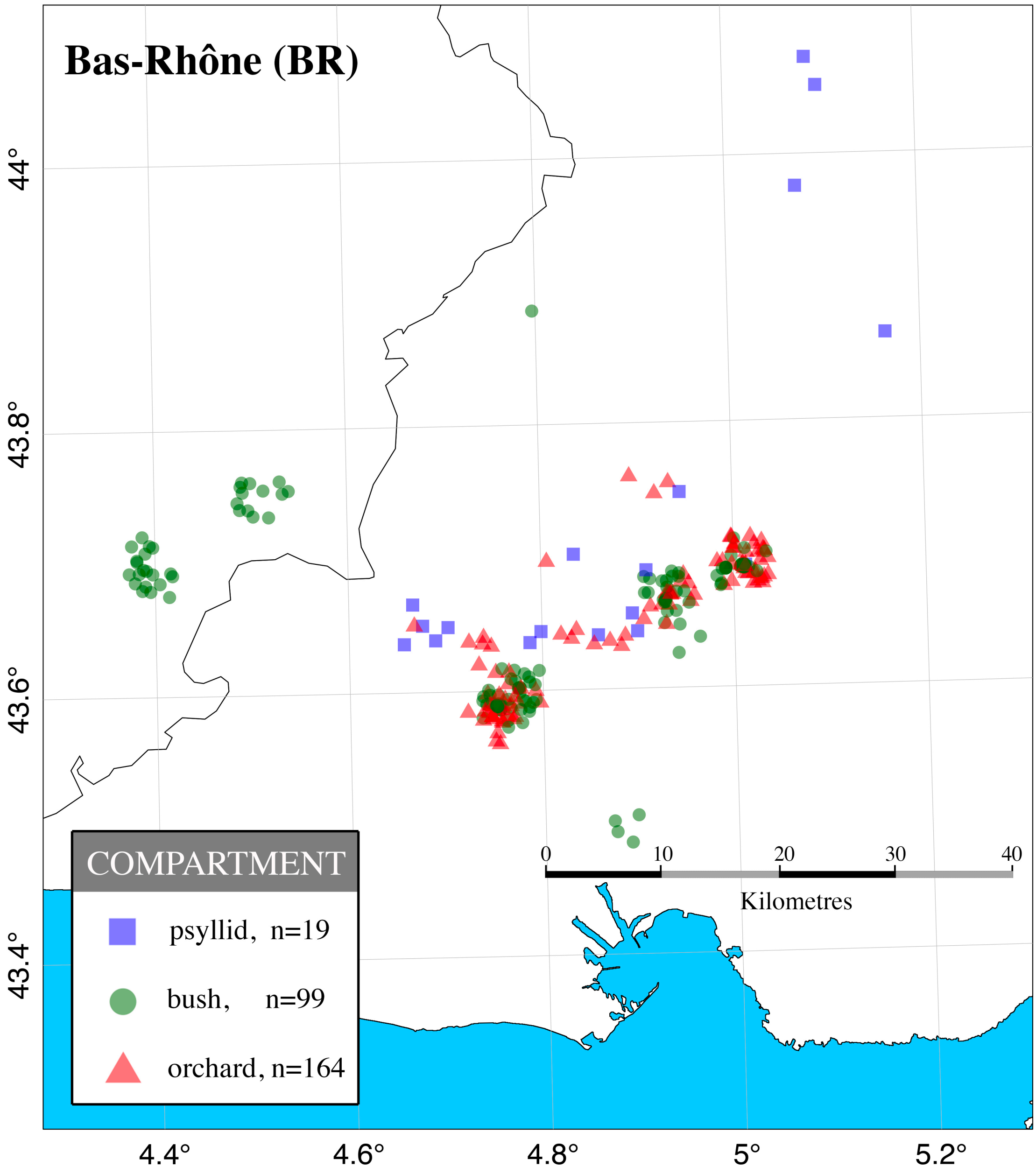

**Figure S5.** Detailed map of the spatial distribution of samples from each compartment (bush, psyllid, and orchard) surveyed in the Valence (VA) region.

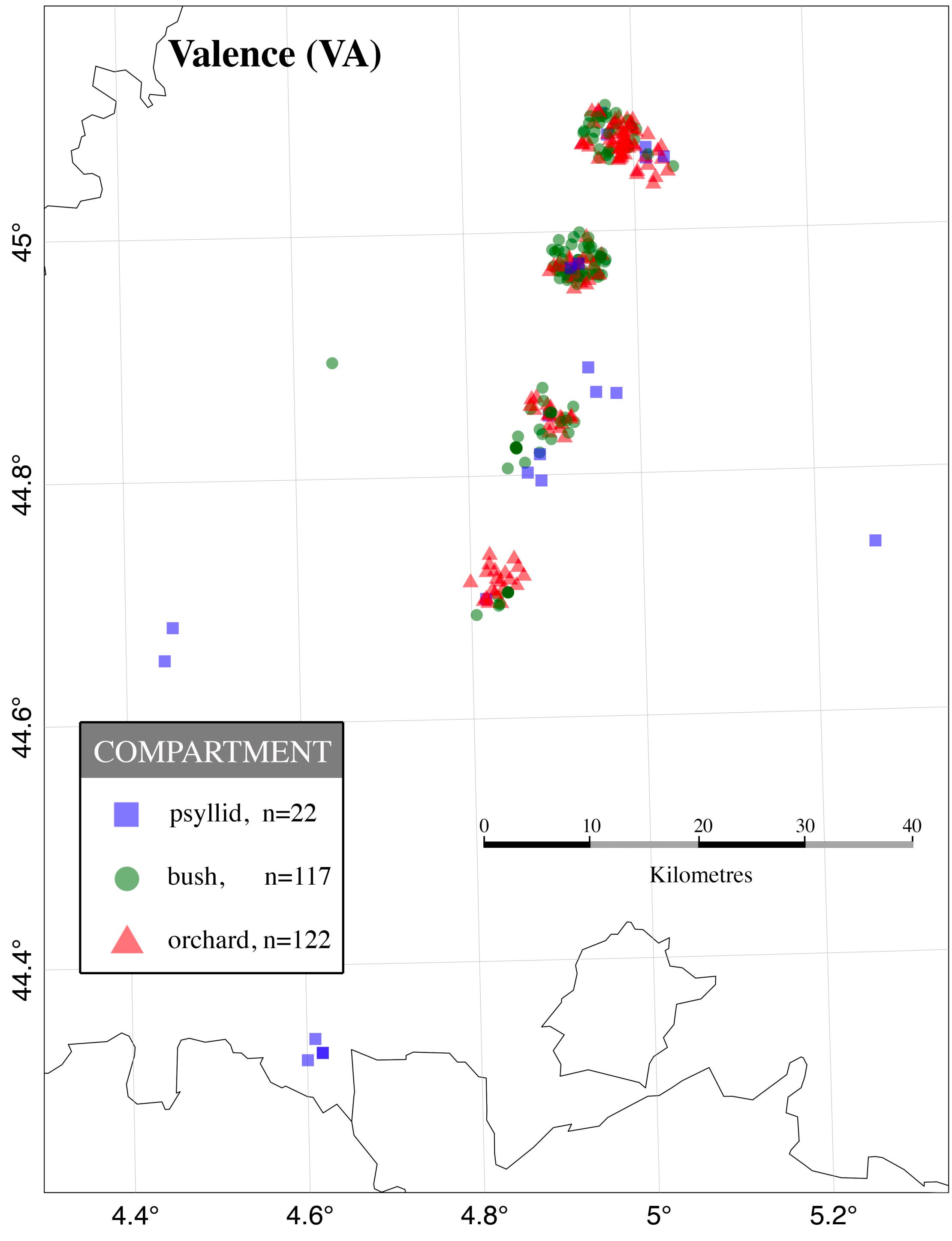

**Supplementary Table S1** Summarized data for all samples surveyed in the study, ordered by region (PO, BR, VA) and ecological compartment (bush, psyllid, orchard).

| Region | Compartment | Host plant | Sub-region | Locality | Bush codename | N | | | #ESFY+ | %ESFY+ | #IMP | Sampling date |
|---|---|---|---|---|---|---|---|---|---|---|---|---|
| PO | bush | wild Prunus | Prades | Bouleternère | M01 | 37 | | | 8 | 21.62 | 6 | 01/04/2010 |
| PO | bush | wild Prunus | Prades | Eus | M02 | 30 | | | 12 | 40.00 | 11 | 01/04/2010 |
| PO | bush | wild Prunus | Prades | Prades | M03 | 46 | | | 36 | 78.26 | 25 | 01/04/2010 |
| PO | bush | wild Prunus | Prades | Sirach | M04 | 12 | | | 3 | 25.00 | 3 | 01/04/2010 |
| PO | bush | wild Prunus | Prades | Codalet | M05 | 16 | | | 8 | 50.00 | 5 | 01/04/2010 |
| PO | bush | wild Prunus | Torreilles | St Laurent de la Salanque | M07 | 26 | | | 4 | 15.38 | 1 | 02/04/2010 |
| PO | bush | wild Prunus | Torreilles | Bompas | M08 | 32 | | | 30 | 93.75 | 21 | 01/04/2010 |
| PO | bush | wild Prunus | Torreilles | Claira | M09 | 25 | | | 0 | 0.00 | 0 | 09/12/2010 |
| PO | bush | wild Prunus | Torreilles | Sica Centrex | M10 | 30 | | | 2 | 6.67 | 1 | 28/04/2011 |
| PO | bush | wild Prunus | Torreilles | Sainte-Marie | M11 | 22 | | | 22 | 100.00 | 18 | 01/04/2010 |
| PO | bush | wild Prunus | Torreilles | Torreilles | M12 | 26 | | | 24 | 92.31 | 19 | 01/04/2010 |
| PO | bush | wild Prunus | Torreilles | Torreilles | M13 | 10 | | | 10 | 100.00 | 9 | 01/04/2010 |
| PO | bush | wild Prunus | Prades | Col de Jau | M90 | 10 | | | 3 | 30.00 | 3 | 12/10/2011 |
| PO | bush | wild Prunus | Prades | Villerach | M91 | 10 | | | 1 | 10.00 | 1 | 10/03/2008 |
| | | | | STATISTICS | N | 14 | | | 14 | 14 | 14 | |
| | | | | | ∑ | 332 | | | 163 | | 123 | |
| | | | | | min | 10 | | | 0 | 0.00 | 0 | |
| | | | | | max | 46 | | | 36 | 100.00 | 25 | |
| | | | | | mean | 23.71 | | | 11.8 | 47.36 | 9.13 | |
| | | | | | SE | 2.96 | | | 3.12 | 10.09 | 2.29 | |
| **Region** | **Compartment** | **Host plant** | **Sub-region** | **Locality** | **Psyllid codename** | **N sp.A+B** | **N sp.A** | **N sp.B** | **#ESFY+** | **%ESFY+** | **#IMP** | **Sampling date** |
| PO | psyllid | wild Prunus | Autre | Valfogona | E025 | 25 | 25 | 0 | 1 | 4.00 | 1 | 23/04/2005 |
| PO | psyllid | wild Prunus | Prades | Col de Jau | E049 | 24 | 0 | 24 | 1 | 4.17 | 1 | 21/11/2005 |
| PO | psyllid | wild Prunus | Prades | Eus | E051 | 27 | 27 | 0 | 1 | 3.70 | 1 | 02/03/2006 |
| PO | psyllid | wild Prunus | Torreilles | Torreilles | E102 | 49 | 29 | 20 | 2 | 4.08 | 2 | ../04/2006 |
| PO | psyllid | wild Prunus | Torreilles | Torreilles | E203 | 77 | 35 | 42 | 5 | 6.49 | 3 | 25/02/2007 |
| PO | psyllid | wild Prunus | Torreilles | Torreilles | E204 | 90 | 45 | 45 | 5 | 5.56 | 4 | 04/03/2007 |
| PO | psyllid | wild Prunus | Torreilles | Torreilles | E205 | 54 | 22 | 32 | 1 | 1.82 | 1 | 11/03/2007 |
| PO | psyllid | wild Prunus | Torreilles | Torreilles | E206 | 30 | 13 | 17 | 2 | 6.67 | 2 | 18/03/2007 |
| PO | psyllid | wild Prunus | Torreilles | Torreilles | E208 | 34 | 9 | 25 | 4 | 11.76 | 4 | 02/04/2007 |
| PO | psyllid | wild Prunus | Torreilles | Torreilles | E212 | 21 | 4 | 17 | 1 | 4.55 | 1 | 23/05/2007 |
| PO | psyllid | wild Prunus | Torreilles | Torreilles | E214 | 7 | 7 | 0 | 1 | 14.28 | 1 | 31/01/2007 |
| PO | psyllid | wild Prunus | Torreilles | Torreilles | E216 | 84 | 48 | 36 | 3 | 3.53 | 3 | 21/02/2007 |
| PO | psyllid | wild Prunus | Torreilles | Torreilles | E217 | 63 | 28 | 35 | 2 | 3.17 | 2 | 28/02/2007 |
| PO | psyllid | wild Prunus | Torreilles | Torreilles | E218 | 108 | 31 | 77 | 2 | 1.80 | 2 | 07/03/2007 |
| PO | psyllid | wild Prunus | Torreilles | Torreilles | E219 | 40 | 13 | 27 | 1 | 2.50 | 1 | 14/03/2007 |
| PO | psyllid | wild Prunus | Torreilles | Torreilles | E220 | 49 | 15 | 34 | 1 | 2.04 | 1 | 28/03/2007 |
| PO | psyllid | wild Prunus | Autre | Capendu | E245 | 28 | 15 | 13 | 2 | 7.14 | 2 | 12/03/2008 |
| PO | psyllid | wild Prunus | Autre | Thézan | E246 | 20 | 12 | 8 | 1 | 5.00 | 1 | 12/03/2008 |
| PO | psyllid | wild Prunus | Prades | Col d'Ares | E255 | 26 | 25 | 1 | 1 | 3.85 | 1 | 14/03/2008 |
| PO | psyllid | wild Prunus | Prades | Palalda | E258 | 30 | 30 | 0 | 1 | 3.13 | 1 | 14/03/2008 |
| PO | psyllid | wild Prunus | Prades | Boule d'Amont | E259 | 48 | 48 | 0 | 2 | 4.17 | 2 | 14/03/2008 |
| PO | psyllid | wild Prunus | Prades | La Bastide R. | E269 | 8 | 3 | 5 | 1 | 12.50 | 1 | 18/03/2008 |
| PO | psyllid | wild Prunus | Prades | Prades | E560 | 104 | 104 | 0 | 2 | 1.92 | 2 | 02/04/2010 |
| PO | psyllid | wild Prunus | Prades | Sirach | E561 | 196 | 189 | 7 | 1 | 0.67 | 1 | 02/04/2010 |
| PO | psyllid | wild Prunus | Prades | Codalet | E562 | 60 | 58 | 2 | 1 | 1.49 | 1 | 02/04/2010 |
| PO | psyllid | wild Prunus | Prades | Rodès | E563 | 82 | 72 | 10 | 1 | 1.06 | 1 | 02/04/2010 |
| PO | psyllid | wild Prunus | Autre | Capestang | E566 | 45 | 6 | 39 | 1 | 2.04 | 1 | 06/04/2010 |
| PO | psyllid | wild Prunus | Autre | Rieux en Minervois | E567 | 42 | 2 | 40 | 2 | 4.00 | 2 | 06/04/2010 |
| PO | psyllid | wild Prunus | Torreilles | St Laurent de la Salanque | E569 | 72 | 32 | 40 | 4 | 5.00 | 4 | 01/04/2010 |
| PO | psyllid | wild Prunus | Torreilles | St Laurent de la Salanque | E665 | 12 | 10 | 2 | 1 | 8.33 | 1 | 02/02/2011 |
| PO | psyllid | wild Prunus | Autre | Puicheric | E744 | 75 | 13 | 62 | 6 | 2.61 | 6 | 28/03/2012 |
| PO | psyllid | wild Prunus | Torreilles | Tautavel | E746 | 93 | 45 | 48 | 1 | 1.07 | 1 | 28/03/2012 |
| | | | | STATISTICS | N | 32 | 32 | 32 | 32 | 32 | 32 | |
| | | | | | ∑ | 1723 | 1015 | 708 | 61 | | 58 | |
| | | | | | min | 7 | 0 | 0 | 1 | 0.67 | 1 | |
| | | | | | max | 196 | 189 | 77 | 6 | 14.29 | 6 | |
| | | | | | mean | 53.84 | 31.72 | 22.13 | 1.91 | 3.54 | 1.81 | |
| | | | | | SE | 6.80 | 6.44 | 3.61 | 0.25 | | 0.22 | |
| **Region** | **Compartment** | **Host plant** | **Sub-region** | **Locality** | **Plot codename** | **N cumulative sampling** | **N per date** | | **#ESFY+** | **%ESFY+** | **#IMP** | **Sampling date** |
| PO | orchard | Prunus armeniaca | Torreilles | Sica Centrex | GL01 | 3 | 3 | | 3 | | 3 | ../04/2008 |
| PO | orchard | Prunus cerasifera | Torreilles | Sica Centrex | GL02 | 1 | 1 | | 1 | | 1 | ../04/2008 |
| PO | orchard | Prunus armeniaca | Torreilles | Sica Centrex | GL03 | 1 | 1 | | 1 | | 1 | ../04/2008 |
| PO | orchard | Prunus armeniaca | Torreilles | Sica Centrex | GL04 | 4 | 4 | | 4 | | 5 | ../04/2008 |
| PO | orchard | Prunus persica | Torreilles | Sica Centrex | GL05 | 2 | 2 | | 2 | | 2 | ../04/2008 |
| PO | orchard | Prunus armeniaca | Torreilles | Marquixanes | GL06 | 2 | 2 | | 2 | | 1 | 03/03/2009 |
| PO | orchard | Prunus armeniaca | Torreilles | Alenya | GL07 | 2 | 2 | | 2 | | 1 | 03/03/2009 |
| PO | orchard | Prunus armeniaca | Torreilles | Villeplaine | GL08 | 1 | 1 | | 1 | | 1 | 03/03/2009 |
| PO | orchard | Prunus armeniaca | Torreilles | Villeneuve | GL09 | 1 | 1 | | 1 | | 1 | 03/03/2009 |
| PO | orchard | Prunus armeniaca | Prades | Ria | P01 | 1 | 1 | | 1 | | 1 | 03/02/2010 |
| PO | orchard | Prunus armeniaca | Prades | Sirach | P02 | 17 | 17 | | 17 | | 17 | 03/02/2010 |
| PO | orchard | Prunus armeniaca | Prades | Codalet | P03a | 3 | 3 | | 3 | | 3 | 03/02/2010 |

| | | | | | | | | | | | | |
|---|---|---|---|---|---|---|---|---|---|---|---|---|
| PO | orchard | Prunus armeniaca | Prades | Codalet | P03b | 118 | 106 | | 26 | 24.56 | 29 | 03/02/2010 |
| PO | orchard | Prunus armeniaca | Prades | Codalet | | | 12 | | 5 | | | 18/03/2011 |
| PO | orchard | Prunus armeniaca | Prades | Lloncet | P04 | 135 | 117 | | 17 | 14.53 | 27 | 03/02/2010 |
| PO | orchard | Prunus armeniaca | Prades | Lloncet | | | 18 | | 18 | | | 10/03/2011 |
| PO | orchard | Prunus armeniaca | Torreilles | Sica Centrex | P05 | 2 | 2 | | 2 | | 2 | 03/02/2010 |
| PO | orchard | Prunus armeniaca | Torreilles | Royal Sophie | P06 | 80 | 2 | | 2 | | 4 | 03/02/2010 |
| PO | orchard | Prunus armeniaca | Torreilles | Royal Sophie | | | 78 | | 4 | 5.13 | | 09/12/2010 |
| PO | orchard | Prunus armeniaca | Torreilles | Claira-Annexe 2 | P07 | 40 | 2 | | 2 | | 2 | 12/02/2008 |
| PO | orchard | Prunus armeniaca | Torreilles | Claira-Annexe 2 | | | 21 | | 21 | | 21 | ../04/2009 |
| PO | orchard | Prunus armeniaca | Torreilles | Claira-Annexe 2 | | | 3 | | 1 | | 0 | 01/04/2010 |
| PO | orchard | Prunus armeniaca | Torreilles | Claira-Annexe 2 | | | 14 | | 14 | | 14 | ../04/2012 |
| PO | orchard | Prunus armeniaca | Torreilles | Claira-Monilia | P08 | 255 | 40 | | 38 | | 81 | 09/12/2010 |
| PO | orchard | Prunus armeniaca | Torreilles | Claira-Monilia | | | 9 | | 8 | | | 05/01/2011 |
| PO | orchard | Prunus armeniaca | Torreilles | Claira-Monilia | | | 206 | | 81 | 39.32 | | 28/04/2011 |
| PO | orchard | Prunus armeniaca | Torreilles | Claira | P10 | 206 | 206 | | 4 | 1.94 | 2 | 09/12/2010 |
| PO | orchard | Prunus armeniaca | Torreilles | Ecoles | P11 | 4 | 4 | | 4 | | 4 | 03/02/2010 |
| PO | orchard | Prunus armeniaca | Torreilles | Entrée Village | P12 | 10 | 10 | | 10 | | 10 | 03/02/2010 |
| PO | orchard | Prunus armeniaca | Torreilles | Sainte-Marie | P13 | 2 | 2 | | 2 | | 1 | 03/02/2010 |
| PO | orchard | Prunus armeniaca | Torreilles | Los Ecorchats | P14 | 3 | 3 | | 3 | | 3 | 03/02/2010 |
| PO | orchard | Prunus armeniaca | Estagel-Rivesaltes | Cases de Pène | P16 | 1 | 1 | | 1 | | 1 | 03/02/2010 |
| PO | orchard | Prunus armeniaca | Estagel-Rivesaltes | Espira de l'Agly | P17 | 2 | 2 | | 2 | | 2 | 17/02/2010 |
| PO | orchard | Prunus armeniaca | Estagel-Rivesaltes | Espira de l'Agly | P18 | 1 | 1 | | 1 | | 1 | 17/02/2010 |
| PO | orchard | Prunus armeniaca | Estagel-Rivesaltes | Rivesaltes | P19 | 1 | 1 | | 1 | | 1 | 17/02/2010 |
| PO | orchard | Prunus armeniaca | Estagel-Rivesaltes | Jonquières | P20 | 4 | 4 | | 4 | | 4 | 17/02/2010 |
| PO | orchard | Prunus armeniaca | Autre | St Genies des Fontaines | P21 | 7 | 7 | | 7 | | 7 | ../04/2009 |
| PO | orchard | Prunus armeniaca | Autre | St Genies des Fontaines | P21 | 5 | 5 | | 5 | | 5 | 17/02/2010 |
| PO | orchard | Prunus armeniaca | Autre | Ceret | P23 | 3 | 3 | | 1 | | 1 | 17/02/2010 |
| PO | orchard | Prunus armeniaca | Autre | Bages | P24 | 2 | 2 | | 2 | | 2 | 17/02/2010 |
| PO | orchard | Prunus armeniaca | Autre | Elne | P26 | 2 | 2 | | 2 | | 2 | 17/02/2010 |
| PO | orchard | Prunus armeniaca | Torreilles | Sica Centrex | P27 | 32 | 32 | | 5 | | 4 | 17/02/2010 |
| | | | | STATISTICS | N | 34 | | | 42 | 5 | 37 | |
| | | | | | ∑ | 953 | | | 331 | | 267 | |
| | | | | | min | 1 | | | 1 | 1.94 | 0 | |
| | | | | | max | 255 | | | 81 | 39.32 | 81 | |
| | | | | | mean | 28.03 | | | 7.88 | 17.09 | 7.22 | |
| | | | | | SE | 10.45 | | | 2.16 | 6.81 | 2.37 | |

| **Region** | **Compartment** | **Host plant** | **Sub-region** | **Locality** | **Bush codename** | **N** | | | **#ESFY+** | **%ESFY+** | **#IMP** | **Sampling date** |
|---|---|---|---|---|---|---|---|---|---|---|---|---|
| BR | bush | wild Prunus | La Dynamite | Arles | M50 | 10 | | | 2 | 20.00 | 1 | 24/03/2009 |
| BR | bush | wild Prunus | La Dynamite | La Dynamite | M51 | 5 | | | 5 | 100.00 | 5 | 24/03/2009 |
| BR | bush | wild Prunus | La Dynamite | La Dynamite | M52 | 20 | | | 10 | 50.00 | 8 | 27/03/2010 |
| BR | bush | wild Prunus | La Dynamite | La Dynamite | M53 | 20 | | | 7 | 35.00 | 6 | 11/04/2010 |
| BR | bush | wild Prunus | La Dynamite | La Dynamite | M54 | 10 | | | 9 | 90.00 | 3 | 16/02/2010 |
| BR | bush | wild Prunus | La Dynamite | La Dynamite | M55 | 10 | | | 2 | 20.00 | 1 | 27/03/2010 |
| BR | bush | wild Prunus | La Samatane | La Samatane | M56 | 15 | | | 1 | 6.67 | 1 | 24/03/2009 |
| BR | bush | wild Prunus | La Samatane | La Samatane | M57 | 10 | | | 2 | 20.00 | 1 | 24/03/2009 |
| BR | bush | wild Prunus | La Samatane | La Samatane | M58 | 10 | | | 8 | 80.00 | 4 | 24/03/2009 |
| BR | bush | wild Prunus | La Samatane | La Samatane | M59 | 10 | | | 2 | 20.00 | 0 | 24/03/2009 |
| BR | bush | wild Prunus | La Samatane | La Samatane | M60 | 20 | | | 3 | 15.00 | 2 | 27/03/2009 |
| BR | bush | wild Prunus | La Samatane | La Samatane | M61 | 10 | | | 2 | 20.00 | 1 | 27/03/2009 |
| BR | bush | wild Prunus | La Samatane | La Samatane | M62 | 10 | | | 2 | 20.00 | 1 | 27/03/2009 |
| BR | bush | wild Prunus | La Samatane | La Samatane | M63 | 10 | | | 3 | 30.00 | 2 | 27/03/2009 |
| BR | bush | wild Prunus | La Samatane | La Samatane | M64 | 10 | | | 1 | 10.00 | 1 | 27/03/2009 |
| BR | bush | wild Prunus | La Samatane | La Samatane | M65 | 10 | | | 4 | 40.00 | 3 | 27/03/2009 |
| BR | bush | wild Prunus | La Samatane | La Samatane | M66 | 10 | | | 5 | 50.00 | 5 | 27/03/2009 |
| BR | bush | wild Prunus | Eyguières | Eyguières | M67 | 10 | | | 5 | 50.00 | 3 | 27/03/2009 |
| BR | bush | wild Prunus | Eyguières | Eyguières | M68 | 20 | | | 8 | 40.00 | 7 | 27/03/2009 |
| BR | bush | wild Prunus | Eyguières | Eyguières | M69 | 10 | | | 5 | 50.00 | 3 | 27/03/2009 |
| BR | bush | wild Prunus | Eyguières | Eyguières | M70 | 10 | | | 8 | 80.00 | 5 | 07/02/2010 |
| BR | bush | wild Prunus | Eyguières | Eyguières | M71 | 20 | | | 6 | 30.00 | 5 | 07/02/2010 |
| BR | bush | wild Prunus | Eyguières | Eyguières | M71 | 20 | | | 6 | 30.00 | | 27/03/2010 |
| BR | bush | wild Prunus | Eyguières | Eyguières | M72 | 10 | | | 2 | 20.00 | 0 | 27/03/2010 |
| BR | bush | wild Prunus | Eyguières | Eyguières | M73 | 10 | | | 9 | 90.00 | 12 | 07/02/2010 |
| BR | bush | wild Prunus | Eyguières | Eyguières | M73 | 20 | | | 5 | 25.00 | | 27/03/2010 |
| BR | bush | wild Prunus | La Samatane | La Samatane | M74 | 10 | | | 2 | 20.00 | 1 | 27/03/2010 |
| BR | bush | wild Prunus | Eyguières | Eygalières | M75 | 10 | | | 4 | 40.00 | 3 | 24/03/2009 |
| BR | bush | wild Prunus | La Dynamite | La Dynamite | M76 | 40 | | | 9 | 22.50 | 6 | 27/03/2010 |
| BR | bush | wild Prunus | La Dynamite | La Dynamite | M77 | 10 | | | 8 | 80.00 | 5 | 11/04/2010 |
| BR | bush | wild Prunus | La Dynamite | La Dynamite | M78 | 6 | | | 3 | 50.00 | 1 | 11/04/2010 |
| BR | bush | wild Prunus | La Dynamite | Saint Martin-de-Crau | M79 | 10 | | | 4 | 40.00 | 3 | 24/03/2009 |
| BR | bush | wild Prunus | La Dynamite | Saint Martin-de-Crau | M80 | 10 | | | 0 | 0.00 | 0 | 24/03/2009 |
| | | | | STATISTICS | N | 33 | | | 33 | 33 | 31 | |
| | | | | | ∑ | 426 | | | 152 | | 99 | |
| | | | | | min | 5 | | | 0 | 0.00 | 0 | |
| | | | | | max | 40 | | | 10 | 100.00 | 12 | |
| | | | | | mean | 12.91 | | | 4.61 | 39.22 | 3.19 | |
| | | | | | SE | 1.15 | | | 0.49 | 4.58 | 0.49 | |

| Region | Compartment | Host plant | Sub-region | Locality | Psyllid codename | N sp.A+B | N sp.A | N sp.B | #ESFY+ | %ESFY+ | #IMP | Sampling date |
|---|---|---|---|---|---|---|---|---|---|---|---|---|
| BR | psyllid | wild Prunus | | Carpentras | E289 | 37 | 36 | 1 | 2 | 5.40 | 2 | 26/03/2008 |
| BR | psyllid | wild Prunus | | Aureilles | E367 | 10 | 10 | 0 | 1 | 10.00 | 1 | 18/03/2009 |
| BR | psyllid | wild Prunus | | Cavaillon | E371 | 58 | 58 | 0 | 2 | 3.45 | 2 | 18/03/2009 |
| BR | psyllid | wild Prunus | | Arles | E380 | 14 | 10 | 4 | 5 | 35.71 | 5 | 21/03/2009 |
| BR | psyllid | wild Prunus | | St Martin de Crau | E381 | 15 | 15 | 0 | 2 | 13.33 | 2 | 21/03/2009 |
| BR | psyllid | wild Prunus | | St Martin de Crau | E383 | 38 | 36 | 2 | 1 | 2.63 | 1 | 21/03/2009 |
| BR | psyllid | wild Prunus | | St Martin de Crau | E385 | 52 | 52 | 0 | 2 | 3.85 | 2 | 21/03/2009 |
| BR | psyllid | wild Prunus | | Eygalières | E386 | 13 | 13 | 0 | 1 | 7.70 | 1 | 21/03/2009 |
| BR | psyllid | wild Prunus | | Mouriès | E387 | 54 | 53 | 1 | 1 | 1.85 | 1 | 21/03/2009 |
| BR | psyllid | wild Prunus | | Peynes-les-Fontaines | E448 | 2 | 2 | 0 | 1 | 50.00 | 1 | 09/03/2009 |
| BR | psyllid | wild Prunus | | Eygalières | E616 | 6 | 6 | 0 | 1 | 16.67 | 1 | 26/03/2006 |
| | | | | STATISTICS | N | 11 | 11 | 11 | 11 | 11 | 11 | |
| | | | | | ∑ | 299 | 291 | 8 | 19 | | 19 | |
| | | | | | min | 2 | 2 | 0 | 1 | 1.85 | 1 | |
| | | | | | max | 58 | 58 | 4 | 5 | 50.00 | 5 | |
| | | | | | mean | 27.18 | 26.46 | 0.73 | 1.73 | 6.35 | 1.73 | |
| | | | | | SE | 6.32 | 6.33 | 0.38 | 0.36 | 4.67 | 0.36 | |

| Region | Compartment | Host plant | Sub-region | Locality | Plot codename | N cumulative sampling | N per date | | #ESFY+ | %ESFY+ | #IMP | Sampling date |
|---|---|---|---|---|---|---|---|---|---|---|---|---|
| BR | orchard | Prunus armeniaca | Saint-Gilles | St Gilles | P60 | 23 | 23 | | 23 | | 18 | ../04/2008 |
| BR | orchard | Prunus persica | Saint-Gilles | Générac | P61 | 2 | 2 | | 2 | | 2 | 09/02/2009 |
| BR | orchard | Prunus domestica | Saint-Gilles | Bellegarde | P62 | 24 | 24 | | 24 | | 13 | ../04/2008 |
| BR | orchard | Prunus salicina | La Dynamite | La Dynamite | P63a | 8 | 8 | | 8 | | 7 | ../04/2002 |
| BR | orchard | Prunus armeniaca | La Dynamite | La Chapelette | P63b | 139 | 13 | | 13 | | 24 | 16/02/2010 |
| BR | orchard | Prunus armeniaca | La Dynamite | La Chapelette | | | 122 | | 20 | 16.39 | | 23/10/2010 |
| BR | orchard | Prunus armeniaca | La Dynamite | La Chapelette | | | 4 | | 2 | | | 26/02/2011 |
| BR | orchard | Prunus armeniaca | La Dynamite | Aulnes | P64a | 140 | 4 | | 4 | | 4 | 16/02/2010 |
| BR | orchard | Prunus armeniaca | La Dynamite | Aulnes | | | 124 | | 14 | 11.29 | 6 | 21/10/2010 |
| BR | orchard | Prunus armeniaca | La Dynamite | Aulnes | | | 12 | | 7 | | 3 | 26/01/2011 |
| BR | orchard | Prunus salicina | La Dynamite | Aulnes | P64b | 23 | 23 | | 8 | | 8 | 26/02/2011 |
| BR | orchard | Prunus armeniaca | Istres | Istres | P65 | 2 | 2 | | 1 | | 1 | ../04/2010 |
| BR | orchard | Prunus armeniaca | Istres | Istres | P66 | 24 | 24 | | 3 | | 3 | ../04/2010 |
| BR | orchard | Prunus armeniaca | La Samatane | La Samatane | P67 | 8 | 8 | | 4 | | 0 | ../04/2006 |
| BR | orchard | Prunus armeniaca | La Samatane | La Samatane | P68a=D11 | 17 | 9 | | 9 | | 12 | ../04/2006 |
| BR | orchard | Prunus armeniaca | La Samatane | La Samatane | | | 8 | | 8 | | | 24/03/2009 |
| BR | orchard | Prunus armeniaca | La Samatane | La Samatane | P68b=D12 | 103 | 101 | | 4 | 3.96 | 3 | 21/10/2010 |
| BR | orchard | Prunus armeniaca | La Samatane | La Samatane | | | 2 | | 1 | | | 26/02/2011 |
| BR | orchard | Prunus armeniaca | La Samatane | La Samatane | P68c=D15 | 118 | 8 | | 8 | | 9 | 24/03/2009 |
| BR | orchard | Prunus armeniaca | La Samatane | La Samatane | | | 110 | | 28 | 25.45 | | 21/10/2010 |
| BR | orchard | Prunus armeniaca | La Samatane | La Samatane | P68d=E15 | 2 | 2 | | 2 | | 2 | 24/03/2009 |
| BR | orchard | Prunus armeniaca | Eyguières | Eyguières | P69 | 8 | 8 | | 8 | | 8 | 16/02/2010 |
| BR | orchard | Prunus armeniaca | Eyguières | Eyguières | P70 | 17 | 17 | | 17 | | 17 | 16/02/2010 |
| BR | orchard | Prunus armeniaca | Eyguières | Eyguières | P71a | 112 | 1 | | 1 | | 1 | 07/02/2010 |
| BR | orchard | Prunus armeniaca | Eyguières | Eyguières | | | 104 | | 5 | 4.81 | 1 | 28/10/2010 |
| BR | orchard | Prunus armeniaca | Eyguières | Eyguières | | | 7 | | 2 | | 1 | 26/02/2011 |
| BR | orchard | Prunus armeniaca | Eyguières | Eyguières | P71b | 119 | 1 | | 1 | | 1 | 07/02/2010 |
| BR | orchard | Prunus armeniaca | Eyguières | Eyguières | | | 117 | | 15 | 12.82 | 5 | 23/10/2010 |
| BR | orchard | Prunus armeniaca | Eyguières | Eyguières | | | 1 | | 0 | | 0 | 26/02/2011 |
| BR | orchard | Prunus armeniaca | Eyguières | Eyguières | P71c | 4 | 1 | | 1 | | 14 | 07/02/2010 |
| BR | orchard | Prunus armeniaca | Eyguières | Eyguières | | | 1 | | 1 | | | 28/10/2010 |
| BR | orchard | Prunus armeniaca | Eyguières | Eyguières | | | 2 | | 2 | | | 26/02/2011 |
| BR | orchard | Prunus armeniaca | Eyguières | Eyguières | P71d | 1 | 1 | | 1 | | | 07/02/2010 |
| BR | orchard | Prunus armeniaca | Eyguières | Eyguières | P71e | 2 | 2 | | 2 | | | 07/02/2010 |
| BR | orchard | Prunus armeniaca | Eyguières | Eyguières | P71f | 2 | 2 | | 2 | | | 27/03/2010 |
| BR | orchard | Prunus armeniaca | Eyguières | Eyguières | P71g | 2 | 2 | | 2 | | | 07/02/2010 |
| BR | orchard | Prunus persica | Avignon | Avignon | P72 | 1 | 1 | | 1 | | 1 | 07/02/2010 |
| | | | | STATISTICS | N | 24 | | | 37 | 6 | 26 | |
| | | | | | ∑ | 901 | | | 254 | | 164 | |
| | | | | | min | 1 | | | 0 | 3.96 | 0 | |
| | | | | | max | 140 | | | 28 | 25.89 | 24 | |
| | | | | | mean | 37.54 | | | 6.86 | 12.45 | 6.31 | |
| | | | | | SE | 10.40 | | | 1.22 | 3.25 | 1.26 | |

| Region | Compartment | Host plant | Sub-region | Locality | Bush codename | N | | | #ESFY+ | %ESFY+ | #IMP | Sampling date |
|---|---|---|---|---|---|---|---|---|---|---|---|---|
| VA | bush | wild Prunus | Romans | Les Balmes | M20 | 31 | | | 22 | 70.97 | 19 | 09/04/2010 |
| VA | bush | wild Prunus | Romans | Saint-Bardoux | M22 | 30 | | | 16 | 53.33 | 16 | 08/04/2010 |
| VA | bush | wild Prunus | Romans | Saint-Bardoux | M23 | 30 | | | 11 | 36.67 | 11 | 09/04/2010 |
| VA | bush | wild Prunus | Romans | Saint-Bardoux | M24 | 33 | | | 12 | 36.36 | 10 | 09/04/2010 |
| VA | bush | wild Prunus | Plaine de Valence | Beauvallon | M25 | 30 | | | 19 | 63.33 | 11 | 09/04/2010 |
| VA | bush | wild Prunus | Plaine de Valence | Beauvallon | M26 | 19 | | | 11 | 57.89 | 6 | 09/04/2010 |
| VA | bush | wild Prunus | Mirmande | Mirmande | M27 | 20 | | | 20 | 100.00 | 6 | 09/04/2010 |
| VA | bush | wild Prunus | Mirmande | Mirmande | M28 | 27 | | | 15 | 55.56 | 13 | 09/04/2010 |
| VA | bush | wild Prunus | Mirmande | Mirmande | M29 | 24 | | | 5 | 20.83 | 1 | 09/04/2010 |
| VA | bush | wild Prunus | Plaine de Valence | Gotheron | M30 | 20 | | | 11 | 55.00 | 8 | 09/04/2010 |

| | | | | | | | | | | | | |
|---|---|---|---|---|---|---|---|---|---|---|---|---|
| VA | bush | wild Prunus | Plaine de Valence | Gotheron | M31 | 18 | | | 0 | 0.00 | 0 | 09/04/2010 |
| VA | bush | wild Prunus | Plaine de Valence | Gotheron | M32 | 16 | | | 1 | 6.25 | 1 | 09/04/2010 |
| VA | bush | wild Prunus | Plaine de Valence | Gotheron | M33 | 26 | | | 1 | 3.84 | 1 | 09/04/2010 |
| VA | bush | wild Prunus | Plaine de Valence | Gotheron | M34 | 32 | | | 15 | 46.87 | 14 | 09/04/2010 |
| | | | | STATISTICS | N | 14 | | | 14 | 14 | 14 | |
| | | | | | ∑ | 356 | | | 159 | | 117 | |
| | | | | | min | 16 | | | 0 | 0.00 | 0 | |
| | | | | | max | 33 | | | 22 | 100.00 | 19 | |
| | | | | | mean | 25.43 | | | 11.36 | 43.35 | 8.36 | |
| | | | | | SE | 1.56 | | | 1.93 | 7.56 | 1.63 | |

| **Region** | **Compartment** | **Host plant** | **Sub-region** | **Locality** | **Psyllid codename** | **N sp.A+B** | **N sp.A** | **N sp.B** | **#ESFY+** | **%ESFY+** | **#IMP** | **Sampling date** |
|---|---|---|---|---|---|---|---|---|---|---|---|---|
| VA | psyllid | wild Prunus | Mirmande | St Marcel d'Ardèche | E003 | 6 | 6 | 0 | 3 | 50.00 | 2 | 12/04/2005 |
| VA | psyllid | wild Prunus | Mirmande | Vesseaux | E094 | 20 | 20 | 0 | 1 | 5.00 | 1 | 13/04/2006 |
| VA | psyllid | wild Prunus | Plaine de Valence | Etoile-sur-Rhône | E117 | 56 | 50 | 6 | 3 | 4.80 | 3 | 15/04/2006 |
| VA | psyllid | wild Prunus | Plaine de Valence | Etoile-sur-Rhône | E119 | 7 | 6 | 1 | 1 | 14.30 | 1 | 09/04/2006 |
| VA | psyllid | wild Prunus | Mirmande | St Marcel d'Ardèche | E321 | 3 | 3 | 0 | 1 | 33.33 | 1 | 29/03/2007 |
| VA | psyllid | wild Prunus | Mirmande | Vesseaux | E441 | 15 | 12 | 3 | 1 | 5.90 | 1 | 14/04/2009 |
| VA | psyllid | wild Prunus | Romans | Saint-Bardoux | E580 | 32 | 7 | 25 | 1 | 2.90 | 1 | 08/04/2010 |
| VA | psyllid | wild Prunus | Romans | Saint-Bardoux | E581 | 32 | 10 | 22 | 1 | 3.26 | 1 | 08/04/2010 |
| VA | psyllid | wild Prunus | Romans | Saint-Bardoux | E582 | 31 | 4 | 27 | 1 | 3.03 | 1 | 09/04/2010 |
| VA | psyllid | wild Prunus | Romans | Saint-Bardoux | E583 | 31 | 9 | 22 | 0 | 0.00 | | 09/04/2010 |
| VA | psyllid | wild Prunus | Romans | Peyrins | E584 | 8 | 2 | 6 | 0 | 0.00 | | 09/04/2010 |
| VA | psyllid | wild Prunus | Romans | Les Balmes | E585 | 45 | 7 | 38 | 3 | 6.25 | 3 | 09/04/2010 |
| VA | psyllid | wild Prunus | Plaine de Valence | Beauvallon | E586 | 76 | 36 | 40 | 3 | 3.26 | 3 | 09/04/2010 |
| VA | psyllid | wild Prunus | Plaine de Valence | Beauvallon | E587 | 25 | 18 | 7 | 0 | 0.00 | | 09/04/2010 |
| VA | psyllid | wild Prunus | Plaine de Valence | Beauvallon | E588 | 11 | 8 | 3 | 1 | 9.09 | 1 | 09/04/2010 |
| VA | psyllid | wild Prunus | Mirmande | Mirmande | E589 | 40 | 36 | 4 | 0 | 0.00 | | 09/04/2010 |
| VA | psyllid | wild Prunus | Mirmande | Mirmande | E590 | 31 | 27 | 4 | 0 | 0.00 | | 09/04/2010 |
| VA | psyllid | wild Prunus | Mirmande | Mirmande | E591 | 1 | 1 | 0 | 1 | 0.00 | 1 | 09/04/2010 |
| VA | psyllid | wild Prunus | Plaine de Valence | Gotheron | E688 | 18 | 14 | 4 | 2 | 8.69 | 2 | 25/03/2011 |
| VA | psyllid | wild Prunus | Plaine de Valence | Gotheron | E689 | 14 | 8 | 6 | 0 | 0.00 | | 24/03/2011 |
| VA | psyllid | wild Prunus | Plaine de Valence | Gotheron | E690 | 2 | 2 | 0 | 0 | 0.00 | | 26/03/2011 |
| VA | psyllid | wild Prunus | Plaine de Valence | Gotheron | E691 | 4 | 3 | 1 | 0 | 0.00 | | 26/03/2011 |
| VA | psyllid | wild Prunus | Plaine de Valence | Gotheron | E692 | 2 | 2 | 0 | 0 | 0.00 | | 26/03/2011 |
| VA | psyllid | wild Prunus | Plaine de Valence | Gotheron | E693 | 1 | 1 | 0 | 0 | 0.00 | | 14/04/2011 |
| VA | psyllid | wild Prunus | Plaine de Valence | Gotheron | E694 | 1 | 1 | 0 | 0 | 0.00 | | 14/04/2011 |
| VA | psyllid | wild Prunus | Plaine de Valence | Gotheron | E695 | 5 | 3 | 2 | 1 | 0.00 | 0 | 14/04/2011 |
| VA | psyllid | wild Prunus | Plaine de Valence | Gotheron | E698 | 10 | 8 | 2 | 0 | 0.00 | | 15/03/2011 |
| VA | psyllid | wild Prunus | Plaine de Valence | Gotheron | E699 | 23 | 1 | 22 | 0 | 0.00 | | 16/06/2011 |
| | | | | STATISTICS | N | 28 | 28 | 28 | 28 | 28 | 15 | |
| | | | | | ∑ | 550 | 305 | 245 | 24 | | 22 | |
| | | | | | min | 1 | 1 | 0 | 0 | 0.00 | 0 | |
| | | | | | max | 76 | 36 | 40 | 3 | 50.00 | 3 | |
| | | | | | mean | 19.64 | 10.89 | 8.75 | 0.86 | 4.36 | 1.47 | |
| | | | | | SE | 3.53 | 2.35 | 2.28 | 0.20 | | 0.24 | |

| **Region** | **Compartment** | **Host plant** | **Sub-region** | **Locality** | **Plot codename** | **N cumulative sampling** | **N per date** | | **#ESFY+** | **%ESFY+** | **#IMP** | **Sampling date** |
|---|---|---|---|---|---|---|---|---|---|---|---|---|
| VA | orchard | Prunus armeniaca | Romans | Romans | P40 | 2 | 2 | | 2 | | 2 | 18/02/2010 |
| VA | orchard | Prunus armeniaca | Romans | Romans | P41 | 102 | 102 | | 0 | | 0 | 18/11/2010 |
| VA | orchard | Prunus armeniaca | Romans | Saint-Bardoux | P42 | 123 | 97 | | 9 | 9.28 | 30 | 18/11/2010 |
| VA | orchard | Prunus armeniaca | Romans | Saint-Bardoux | | | 26 | | 26 | | | 15/03/2011 |
| VA | orchard | Prunus armeniaca | Plaine de Valence | Beauvallon | P43 | 111 | 5 | | 5 | | 17 | 18/02/2010 |
| VA | orchard | Prunus armeniaca | Plaine de Valence | Beauvallon | | | 100 | | 6 | 6.00 | | 18/11/2010 |
| VA | orchard | Prunus armeniaca | Plaine de Valence | Beauvallon | | | 6 | | 6 | | | 15/03/2011 |
| VA | orchard | Prunus armeniaca | Plaine de Valence | Etoile-sur-Rhône | P44 | 6 | 6 | | 6 | | 6 | 18/02/2010 |
| VA | orchard | Prunus armeniaca | Mirmande | Mirmande | P45 | 8 | 8 | | 8 | | 8 | 18/02/2010 |
| VA | orchard | Prunus armeniaca | Mirmande | Mirmande | P46 | 2 | 2 | | 2 | | 1 | 18/02/2010 |
| VA | orchard | Prunus armeniaca | Mirmande | Mirmande | P47 | 152 | 150 | | 4 | 2.67 | 3 | 19/10/2011 |
| VA | orchard | Prunus armeniaca | Mirmande | Mirmande | | | 2 | | 0 | | | 15/03/2011 |
| VA | orchard | Prunus armeniaca | Mirmande | Villes-sur-Auzon | P48 | 14 | 14 | | 14 | | 0 | 18/02/2010 |
| VA | orchard | Prunus armeniaca | Plaine de Valence | Gotheron | P49 | 279 | 8 | | 8 | | 52 | 26/10/2010 |
| VA | orchard | Prunus armeniaca | Plaine de Valence | Gotheron | | | 14 | | 14 | | | 26/10/2010 |
| VA | orchard | Prunus armeniaca | Plaine de Valence | Gotheron | | | 244 | | 47 | 19.26 | | 29/03/2011 |
| VA | orchard | Prunus armeniaca | Plaine de Valence | Gotheron | | | 13 | | 5 | | | 16/06/2011 |
| VA | orchard | Prunus armeniaca | Plaine de Valence | Etoile-sur-Rhône | P50 | 3 | 3 | | 3 | | 3 | 03/03/2009 |
| | | | | STATISTICS | N | 11 | | | 18 | 4 | 11 | |
| | | | | | ∑ | 802 | | | 165 | | 122 | |
| | | | | | min | 2 | | | 0 | 2.67 | 0 | |
| | | | | | max | 279 | | | 47 | 19.26 | 52 | |
| | | | | | mean | 72.91 | | | 9.17 | 9.30 | 11.09 | |
| | | | | | SE | 27.08 | | | 2.66 | 3.58 | 4.92 | |

**Supplementary Table S2 -** Contingency table of the frequency distribution of all haplotypes successfully genotyped in each ecological compartment (bush, psyllid, and orchard) and each of the three growing regions.

| Haplotype | PO | | | BR | | | VA | | | | |
|---|---|---|---|---|---|---|---|---|---|---|---|
| | Bush | Psyllid | Orchard | Bush | Psyllid | Orchard | Bush | Psyllid | Orchard | ∑ | % |
| I01 | 46 | 28 | 128 | 8 | 5 | 82 | 61 | 6 | 74 | 438 | 44.2 |
| *I01-104** | | | 5 | | | | 1 | | | 6 | 0.6 |
| *I01-248** | | | | | | | | | 1 | 1 | 0.1 |
| *I01-339* | | | | | 1 | | | | | 1 | 0.1 |
| I04 | 5 | 4 | 17 | 6 | 1 | 13 | 2 | 1 | 5 | 54 | 5.4 |
| *I04-09* | | | | | | | | | 1 | 1 | 0.1 |
| *I04-175** | | | | | | 1 | | | | 1 | 0.1 |
| *I04-407** | | | | | | 1 | | | | 1 | 0.1 |
| *I04-453* | | | | | | 1 | | | | 1 | 0.1 |
| I03 | | | | | | | 1 | | | 1 | 0.1 |
| I09 | 37 | 8 | 55 | 71 | 7 | 34 | 48 | 12 | 25 | 297 | 30.0 |
| I10 | 6 | 3 | 8 | 5 | | 15 | 1 | 3 | 15 | 56 | 5.7 |
| *I10-307** | | | 1 | | | | | | | 1 | 0.1 |
| I10-310* | | | 1 | | | | | | | 1 | 0.1 |
| I11 | 25 | 12 | 32 | 6 | 1 | 3 | 3 | | | 82 | 8.3 |
| *I11-267* | | | 1 | | | | | | | 1 | 0.1 |
| I13 | 4 | 3 | 19 | 3 | 4 | 14 | | | 1 | 48 | 4.8 |
| | 123 | 58 | 267 | 99 | 19 | 164 | 117 | 23 | 122 | 991 | 100 |

**Supplementary Table S3 -** Contingency table displaying the frequency distribution of the six major *imp* haplotypes in each ecological compartment (bush, psyllid, and orchard) and each of the three growing regions of the study.

| Haplotype | PO | | | BR | | | VA | | | |
|---|---|---|---|---|---|---|---|---|---|---|
| | Bush | Psyllid | Orchard | Bush | Psyllid | Orchard | Bush | Psyllid | Orchard | ∑ |
| I01 | 46 | 28 | 128 | 8 | 5 | 82 | 61 | 6 | 74 | 438 |
| I04 | 5 | 4 | 17 | 6 | 1 | 13 | 2 | 1 | 5 | 54 |
| I09 | 37 | 8 | 55 | 71 | 7 | 34 | 48 | 12 | 25 | 297 |
| I10 | 6 | 3 | 8 | 5 | 0 | 15 | 1 | 3 | 15 | 56 |
| I11 | 25 | 12 | 32 | 6 | 1 | 3 | 3 | 0 | 0 | 82 |
| I13 | 4 | 3 | 19 | 3 | 4 | 14 | 0 | 0 | 1 | 48 |
| ∑ | 123 | 58 | 259 | 99 | 18 | 161 | 115 | 22 | 120 | 975 |

**Supplementary Table S4 -** Relative proportions of the six major haplotypes in the three ecological compartments and the three regions.

| Haplotype | PO | | | BR | | | VA | | |
|---|---|---|---|---|---|---|---|---|---|
| | Bush | Psyllid | Orchard | Bush | Psyllid | Orchard | Bush | Psyllid | Orchard |
| I01 | 37.4 | 48.3 | 49.4 | 8.1 | 27.8 | 50.9 | 53.0 | 27.3 | 61.7 |
| I04 | 4.1 | 6.9 | 6.6 | 6.1 | 5.6 | 8.1 | 1.7 | 4.5 | 4.2 |
| I09 | 30.1 | 13.8 | 21.2 | 71.7 | 38.9 | 21.1 | 41.7 | 54.5 | 20.8 |
| I10 | 4.9 | 5.2 | 3.1 | 5.1 | 0.0 | 9.3 | 0.9 | 13.6 | 12.5 |
| I11 | 20.3 | 20.7 | 12.4 | 6.1 | 5.6 | 1.9 | 2.6 | 0.0 | 0.0 |
| I13 | 3.3 | 5.2 | 7.3 | 3.0 | 22.2 | 8.7 | 0.0 | 0.0 | 0.8 |
| ∑ | 100 | 100 | 100 | 100 | 100 | 100 | 100 | 100 | 100 |

**Supplementary Table S5 -** Number of major *imp* haplotypes genotyped for each species of the *Cacopsylla pruni* complex, all regions combined, and region by region.

| Region | Haplotype | Species A | Species B |
|---|---|---|---|
| PO+BR+VA | I01 | 12 | 27 |
| | I04 | 4 | 2 |
| | I09 | 22 | 5 |
| | I10 | 5 | 1 |
| | I11 | 10 | 3 |
| | I13 | 5 | 2 |
| | **#A/B** | **1611** | **961** |
| | **#ESFY+** | **58** | **40** |
| | **%A/B** | **62.6** | **37.4** |
| | **%ESFY+** | **59.2** | **40.8** |
| PO | I01 | 7 | 21 |
| | I04 | 2 | 2 |
| | I09 | 5 | 3 |
| | I10 | 2 | 1 |
| | I11 | 9 | 3 |
| | I13 | 2 | 1 |
| | **#A/B** | **1015** | **708** |
| | **#ESFY+** | **27** | **31** |
| | **%A/B** | **58.9** | **41.1** |
| | **%ESFY+** | **46.6** | **53.4** |
| BR | I01 | 3 | 2 |
| | I04 | 1 | 0 |
| | I09 | 7 | 0 |
| | I10 | 0 | 0 |
| | I11 | 1 | 0 |
| | I13 | 3 | 1 |
| | **#A/B** | **291** | **8** |
| | **#ESFY+** | **15** | **3** |
| | **%A/B** | **97.3** | **2.7** |
| | **%ESFY+** | **83.3** | **16.7** |
| VA | I01 | 2 | 4 |
| | I04 | 1 | 0 |
| | I09 | 10 | 2 |
| | I10 | 3 | 0 |
| | I11 | 0 | 0 |
| | I13 | 0 | 0 |
| | **#A/B** | **305** | **245** |
| | **#ESFY+** | **16** | **6** |
| | **%A/B** | **55.5** | **44.5** |
| | **%ESFY+** | **72.7** | **27.3** |

**Supplementary Table S6** List of the *imp* haplotypes described to date, with their GenBank accession numbers.

| *imp* genotype | Genbank Accession n° | Reference |
|---|---|---|
| I01 | FN600707 | Danet et al., 2011 |
| I01-104 | MN116709 | this study |
| I01-248 | MN116710 | this study |
| I01-339 | MN116711 | this study |
| I01-409 | MN116712 | this study |
| I02 | FN600708 | Danet et al., 2011 |
| I03 | FN600709 | Danet et al., 2011 |
| I04 | FN600710 | Danet et al., 2011 |
| I04-9 | MN116713 | this study |
| I04-175 | MN116714 | this study |
| I04-407 | MN116715 | this study |
| I04-453 | MN116716 | this study |
| I05 | FN600711 | Danet et al., 2011 |
| I06 | FN600712 | Danet et al., 2011 |
| I07 | FN600713 | Danet et al., 2011 |
| I08 | FN600714 | Danet et al., 2011 |
| I09 | FN600715 | Danet et al., 2011 |
| I10 | FN600716 | Danet et al., 2011 |
| I10-307 | MN116717 | this study |
| I10-310 (=I34) | MG972433 | Dermastia et al., 2018 |
| I11 | FN600717 | Danet et al., 2011 |
| I11-267 | MN116718 | this study |
| I12 | FN600718 | Danet et al., 2011 |
| I13 | FN600719 | Danet et al., 2011 |
| I26 | FN600720 | Danet et al., 2011 |
